# The Adhesion GPCR Flamingo-Like 1 (FMIL-1) Directs Synapse Formation in a Nociceptive Circuit

**DOI:** 10.64898/2026.09.01.748526

**Authors:** Tyler Kennedy, Damilola Oje, Kylie Howerter, Susan Reese, Rebecca McWhirter, Barbara M.J. O’Brien, Jamie Stern, Morgan Ottley, Demet Araç, Engin Özkan, Richard Sando, Andrew Dillin, David M. Miller

## Abstract

The species-specific anatomy of nervous systems suggests that circuit architecture is largely encoded by genetic blueprints. In *C. elegans,* PVD nociceptive neurons synapse with PVC and AVA interneurons to drive escape responses to noxious stimuli. We used fluorescent markers for PVD synapses with PVC and AVA in a candidate screen to detect connectivity genes. This approach revealed that the LIM homeodomain transcription factor MEC-3 and its target, FMIL-1 (Flamingo-like), function in PVD to direct connectivity with PVC and AVA. FMIL-1 is an adhesion class G Protein-coupled receptor (aGPCR), a protein family with members also implicated in mammalian synapse formation. We show that FMIL-1 acts early in PVD and is also sufficient to induce ectopic synapses in another circuit, thus suggesting that FMIL-1 promotes synaptogenesis. Our work establishes a new experimental circuit in *C. elegans* for investigating neuronal connectivity and provides evidence that FMIL-1/aGPCRs regulate synapse formation.

**Teaser:** FMIL-1 is a synaptogenic adhesion GPCR protein homologous to vertebrate proteins with similar roles.

## Introduction

The nervous system can react to noxious stimuli by triggering motor circuit dependent escape responses. This sensorimotor axis is defined by stereotypical patterns of connectivity that link sensory receptors to the spinal cord and brain. In nervous systems mapped at synaptic resolution, such as *C. elegans,* the neuron specificity of synapses is preserved among genetically identical animals (*1–3*). Notably, despite the densely packed *C. elegans* neuropile, connectivity is limited to a subset of the neurons that contact one another, which argues for the existence of specific synaptogenic determinants (*4–7*).

Members of conserved secreted and cell adhesion proteins have been found to direct synaptic specificity in *C. elegans*, including *unc-6*/Netrin-1 and its receptor, *unc-40/*DCC, *nrx-1*/Neurexin, *lin-17*/Frizzled, *syg-1*/Kirrel3 and *casy-1*/Calsyntenin (*8–17*). The limited number of synapses to which these cell surface proteins have been assigned suggests that multiple additional determinants of synaptogenesis have yet to be identified (*5–7*). As in *C. elegans,* vertebrate synapses may also depend on unique interacting proteins that govern the selection of synaptic partners (*14*, *18*). One implication of this broad deployment of target selection proteins is that genetic mutations in these molecules appear to affect small but cumulative impacts on brain development. For example, genes implicated in target selection are highly represented in genome wide association studies of disorders such as autism spectrum disorder (ASD) and intellectual disability (ID) (*19*). Severity is often linked to accumulated mutations, suggesting additive disruptions to developing circuits (*20*, *21*). Thus, identifying these genes and determining their modes of action is critical for ameliorating brain neuropathologies (*22*).

Here we investigated genetic determinants of glutamatergic synapses between the PVD nociceptive neuron and its post-synaptic partners, the PVC and AVA interneurons in *C. elegans* (*23*, *24*). A pair of PVD neurons, each on opposite sides of the body, arises during postembryonic development in the second larval (L2) stage. Each PVD neuron adopts a stereotypically branched dendritic arbor immediately beneath the epidermis that responds to polymodal inputs, including harsh touch, cold temperature, hyperosmolarity and sound (*25–28*). A single axon from each PVD neuron synapses with PVC and AVA to evoke an escape response (*24*, *29*, *30*). Despite the densely packed VNC neuropile, PVD synaptic targets are limited to PVC and AVA (*3*). Thus, assembly of this simple circuit likely depends on genetic determinants that regulate synaptic specificity(*2*, *31*). Importantly, due to its creation during larval development, the PVD circuit is readily accessible to live imaging and genetic analysis (*32–34*).

Here we show that MEC-3, a LIM homeodomain transcription factor that directs PVD dendritic branching, also promotes the creation of PVD synapses with PVC and AVA (*35*). From a candidate genetic screen of MEC-3-regulated transcripts, we determined that the Flamingo-Like-1 (FMIL-1) protein functions in PVD to induce synapse formation with PVC and AVA. FMIL-1 is a member of the adhesion GPCR (aGPCR) family of proteins. The most closely related vertebrate homologs, the Cadherin EGF LAG seven-pass G-type receptors (CELSRs), are also synaptogenic and have been linked to epilepsy and other neurological disorders (*36–40*). aGPCRs are characterized by a large extracellular region with multiple classes of adhesion domains, as well as a self-cleaving GAIN (GPCR Autoproteolysis-INducing) domain. We show that the synaptogenic role of FMIL-1 depends on its extracellular adhesion domains, but that cleavage of the GAIN domain is not required. Importantly, we demonstrate that neuron-specific expression of FMIL-1 is sufficient to induce ectopic synapses in another circuit. We present a new model of synaptogenesis in which an aGPCR is expressed in pre-synaptic neurons to induce synapse formation.

## Methods

### Strains

*C. elegans* strains (Sup table 1) were grown at 20°C on OP50-1 *Escherichia coli-*seeded nematode growth medium (NGM) plates (*41*). For RNA-seq experiments, *C. elegans* strains were grown on 8P nutrient agar seeded with *E. coli* strain NA22.

### Molecular Biology

We used either the In-Fusion (Takara) or NEBuilder (NEB) cloning kit for all plasmids generated in this study. The *fmil-1a.1* isoform cDNA was cloned using RT-PCR with Protoscript II (NEB). The *nlp-82* promoter was generated by cloning a 4kb region upstream of the *nlp-82* start codon. Isoforms and deletions used for FMIL-1 rescue experiments are listed in Sup Table 1. Plasmids were injected using standard methods (*42*). The *wdIs128*(*F49H12.4::Chrimson)* line was generated using x-ray irradiation and outcrossed at least 7 times (*43*). The *fmil-1::spGFP11x7* was generated by CRISPR insertion by SUNY Biotech. The *fmil-1(T655G) and (H657G)* strains were generated by CRISPR site-directed mutagenesis with a 90bp repair template and selected using a *dpy-5* co-CRISPR (*44*). PCR-verified CRISPR strains were outcrossed 6 times. All mutant strains and crosses were verified by PCR and/or Sanger sequencing.

### Imaging

Images were collected on either a Nikon A1R or a Zeiss 900 confocal microscope with a 40x objective. Animals were mounted on a 5% agar pad, immobilized with 0.025% Tricaine, 9mM Levamisole, and 50mM Muscimol (*45*, *46*). Coverslips were sealed with a 50:50 mixture of Vaseline (Unilever) and paraplast (Sigma-Aldrich) (*33*). Animal age was either staged by hour post lay (HPL) after a 1-hour laying period or by assessing the developmental stage of the vulva for L4 larvae (*47*). Young adults were identified as animals with complete vulval development but no visible eggs.

### Image Analysis

Image analysis was performed using FIJI (*48*). Collected images were flattened using max z projection. A 5-pixel wide line was traced along the visible PVD GFP::RAB-3 puncta. The average value along the line for collected channels (GFP and scarlet for PVD-PVC synapses) was plotted in a Microsoft Excel spreadsheet. Each single and co-localized peak was quantified between the beginning of the PVD axon and the vulva. Axon length was calculated by measuring the length from the first synaptic puncta to the last, or the vulva, whichever came first. All images were blinded by randomizing the file names prior to analysis.

### Live image capture and analysis

Live developmental images were captured on a Nikon SoRa spinning disk confocal microscope. Animals were synchronized by collecting embryos from hypochlorite-treated adult hermaphrodites and allowing them to hatch overnight on unseeded plates at 20°C . 22-24 hours after the hypochlorite treatment, the stalled L1 animals were transferred to OP50 seeded plates to resume development. After 20 hours of feeding, animals were at approximately 34-36hpl and mounted as noted in the *Imaging* section, using S-basal as the supporting medium (*49*). Animals were imaged with a 40x objective for 8 hours with image collection every 3 minutes (161 frames). Imaging files were converted to max Z projections. Using a custom FIJI script based on the “Batch Axon Tracer – sld” macro in the wormSNAP package, axons were straightened through the time series and then processed with wormSNAP to quantify GFP::RAB-3 puncta across time (*50*). We further aligned the straightened images using the Translate and Template Matching functions in FIJI to generate movies and montage images of RAB-3 appearance (*48*, *51*).

### Isolation of PVD neurons by Fluorescence Activated Cell Sorting (FACS)

We used genetic crosses with available fluorescent reporter lines (*otIs181, otIs138, otIs396*) to construct a multicolor marker strain *NC3182* in which PVD is uniquely labeled with GFP. (Sup Fig 1A) (*52*). Wild-type and *mec-3* mutant *NC3182* animals were synchronized and grown to the late L2/early L3 stage at 20°C on 150mm 8P plates seeded with the *E. coli* strain NA22. Larval cells were dissociated as previously described and incubated with DAPI to label damaged cells (*53*, *54*). Larval cells were also isolated from the N2 strain and from *otIs138* to set FACS gates for auto-fluorescent and GFP^+^ cells, respectively. We used a BD FACS Aria in the Vanderbilt Flow Cytometry Core to isolate GFP+ cells (PVD neurons) from the *NC3182* preparation of dissociated cells. GFP^+^ cells were sorted directly into Trizol-LS (Invitrogen) and stored at -80°C. For a preparation of a whole animal reference RNA, aliquots of L2/L3 larval wild-type and *mec-3* NC3182 animals were flash-frozen in liquid N2 prior to the cell dissociation procedure. Triplicate samples were obtained for each genotype for dissociated cells and for whole animal references (12 total samples).

**Figure 1:**
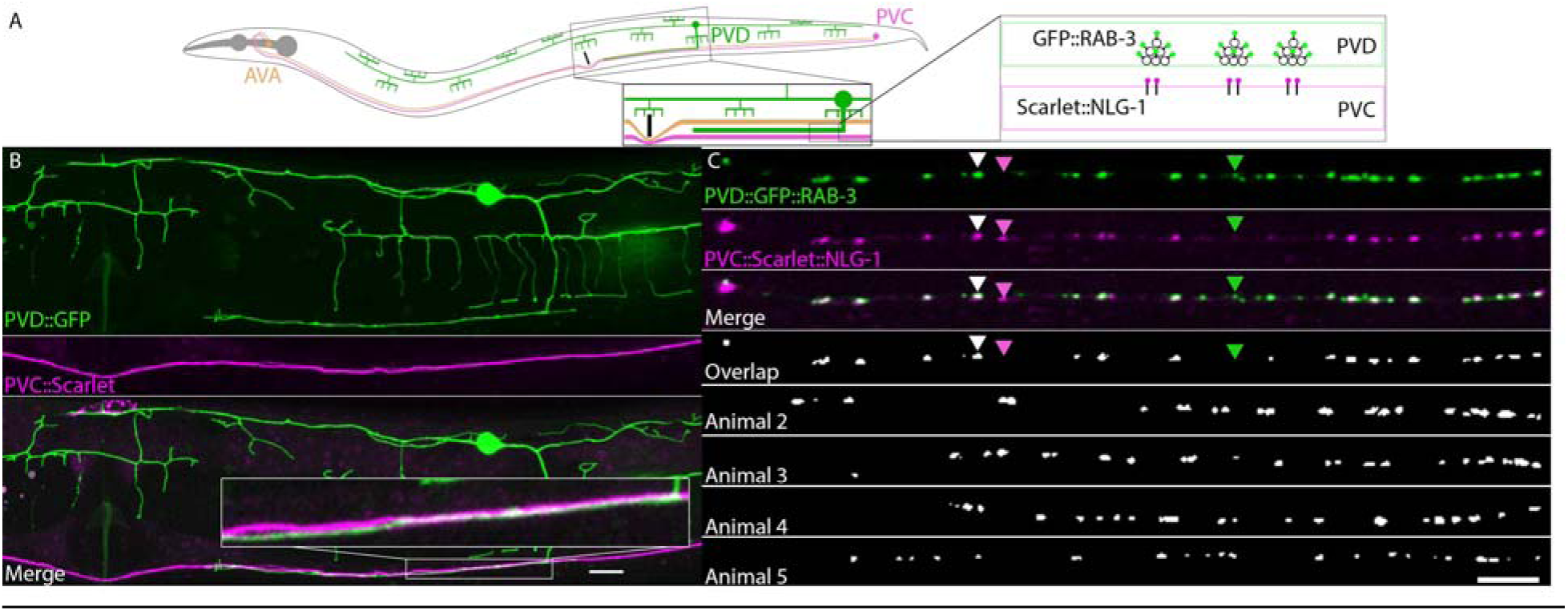
The PVD nociceptive neurons synapse with specific interneuron targets, PVC and AVA, in the ventral nerve cord. A. Morphology and positions of the PVD nociceptive neuron and its interneuron targets, PVC and AVA. (Left) The PVD axon fasciculates with PVC and AVA in the ventral nerve cord (VNC), posterior to the vulva (vertical black line). (Right) Labeling strategy to observe PVD synapses with PVC. GFP::RAB-3 labels PVD pre-synaptic clusters of synaptic vesicles and Scarlet::NLG-1/Neuroligin labels PVC post-synaptic domains. B. Neuron-specific markers PVD::GFP (green) (*Poig-8::GFP)*, and PVC::Scarlet (magenta) (*Pnlp-82*::Scarlet). Scale bar 10 µm. Inset: PVD axon and PVC neurite tightly fasciculate. C. Co-localization of PVD pre-synaptic domains (GFP::RAB-3) with PVC post-synapses (Scarlet::NLG-1) (Merge). Overlap of RAB-3/NLG-1 puncta (white) is shown for 5 representative animals. Examples of overlapping (white arrowhead) and rare “unpaired” pre-synaptic GFP::RAB-3 puncta (green arrowhead) and post-synaptic Scarlet::NLG-1 (red arrowhead) are indicated. Scale bar 10 µm.

### Bulk RNA-sequencing of FACS-isolated cells and reference samples

Frozen pellets of whole L2/L3 larval stage reference samples (N2 and *mec-3 NC3182*) were ground to powder in a mortar and pestle chilled with liquid nitrogen and dissolved in Trizol-LS. RNA was extracted from Trizol-LS preparations of dissociated cells and whole animal references and DNA contamination removed using the Zymo DNA-free RNA Kit (Zymo Research, Irving, CA) according to manufacturer’s instructions. RNA quality was determined by an Agilent Bioanalyzer at the VANTAGE Core. Samples with RNA Integrity Numbers (RIN) scores of ≥ 7 and 5-10 ng of total RNA were converted to cDNA using poly-dT primers to capture mRNA and amplified prior to library preparation using Takara-Clontech SMARTer technology. Libraries were constructed using an Illumina kit and sequenced on an Illumina HiSeq 3000 system to generate 75 bp paired-end reads (30-60 million reads per sample).

### Analysis of RNA-seq data

Differentially expressed transcripts were obtained using the RNA-Seq Differential Expression analysis pipeline in CLC Genomics Workbench which utilizes a negative binomial GLM model. Transcripts were scored as differentially expressed >2-fold difference and FDR-corrected p value <0.05 compared to control (Fig 2C, Sup Table 2, 4). For our initial study of PVD we focused on transcription factors (TFs) as likely to produce strong phenotypes due to regulation of multiple downstream genes (*55–59*). We identified a total of 63 TFs below a 1% FDR cutoff, with at least 10TPM and enriched at least 2-fold in PVD against all other cell types (*60*). We also examined the CeNGEN L4 dataset and identified 76 TFs with at least 10TPM expression, enriched at least 2-fold in PVD over all other neurons (*61*). 23 TFs overlapped across the CeNGEN and L2/L3 bulk RNA-seq dataset (*61*) (Sup Table 3). Once we identified a *mec-3* synaptic phenotype we explored its transcriptional regulation. *mec-3* is typically considered a transcriptional activator and accordingly most targets below a 1% FDR (260) were downregulated in *mec-3* mutants, while the remainder (58) were upregulated (*62*). We focused on the seven downregulated genes described in the results based on protein domain annotations associated with synapse formation and relative enrichment in PVD neurons vs all neurons in the L4 stage CeNGEN dataset.

**Figure 2:**
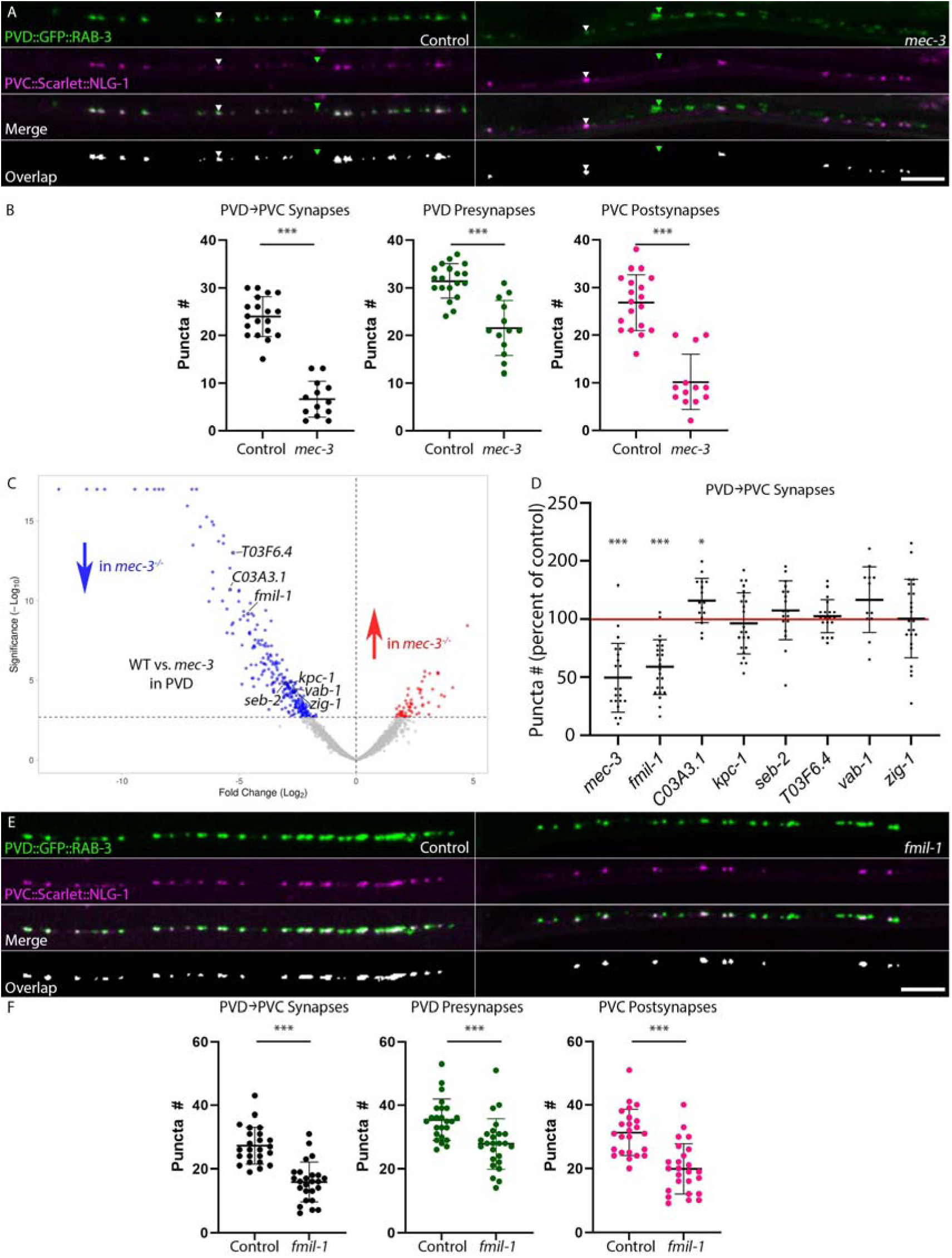
The LIM-homeodomain transcription factor MEC-3 and its downstream target FMIL-1 promote the formation of PVD→PVC synapses. A. PVD pre-synapses (GFP::RAB-3) and PVC post-synapses (Scarlet::NLG-1) are reduced in *mec-3(gk1126)* mutants relative to control, resulting in fewer PVD→PVC synapses (Merge and Overlap). Overlapping PVD→PVC synaptic puncta (white) are depicted in the bottom panel (Overlap). Examples of overlapping (white arrowhead) and rare “unpaired” pre-synaptic GFP::RAB-3 puncta (green arrowhead) are indicated. Scale bar 10µm. B. Quantification of total PVD→PVC synapses (p = 6.7x10^-13^), total pre-synaptic number (p = 1.5x10^-6^) and total post-synaptic number (p = 7.4x10^-9^) between control and *mec-3(gk1126)* null mutants. n≥13 for all experiments. Error bars are SD. Student’s t-test. C. mRNA fold change volcano plot for PVD transcripts in control vs. *mec-3* mutant animals. PVD transcripts down-regulated (blue arrow) vs upregulated (red) in *mec-3* mutants. Down-regulated genes of interest are labeled. Significance thresholds are ≥2x fold difference, 1% FDR (dashed horizontal line). D. Quantification of PVD→PVC synaptic number in mutants of *mec-3* and downstream genes vs control. Fewer PVD→PVC synapses were observed for *mec-3(e1338)* (p=4.8x10^-5^) and *fmil-1(gk643828)* (p=9.8x10^-7^). All other alleles are NS. n≥10 for all experiments. Error bars represent SD. Student’s t-test. E. PVD pre-synapses (GFP::RAB-3) and PVC post-synapses (Scarlet::NLG-1) are reduced in *fmil-1(ok2115)* mutant animals vs control, resulting in fewer PVD→PVC synapses (Merge and Overlap). Scale bar 10µm. F. Quantification of PVD→PVC synapses (p=3.0x10^-8^), pre-synapse number (p=6.9x10^-4^) and post-synapse number (p=3.3x10^-6^) between control and *fmil-1(ok2115)* nulls. n≥24 for all experiments. Error bars represent SD. Student’s t-test.

### RNAi treatment

Animals in the *eri-1* background, expressing *ser-2prom-3p*::GFP::RAB-3 and *nlp-82p*::Scarlet::NLG-1 were grown at 20 ° C on bacterial lawns expressing dsRNA of interest or an empty vector (EV) prepared using previously established techniques (*63*). Adults were allowed to lay for 1 day and then removed. The resulting offspring were imaged 2 days later at the L4 stage.

### Protein Domain Analysis

Protein domains for FMIL-1 were inferred using AlphaFold Protein Structure Database entry AF-U4PMB9-F1-v4 by visual inspection and by structure searches using Foldseek against known structures and the CATH database, sequence searches against the SMART database and sequence alignments with other adhesion GPCRs (*64–68*). We identified two N-terminal domains belonging unambiguously to the Cadherin superfamily. The third domain has an Ig-like topology but lacks some features of classical Ig domains and may be classified as a Fibronectin-type III domain based on sequence searches. The remaining domains are EGF-like, Hormone-binding domain and GAIN domains, followed by a seven-pass transmembrane domain.

*in silico* tethered agonist (TA) binding was assessed using ColabFold (*69*). *C. elegans* aGPCRs were truncated at their predicted cleavage site and analyzed using the default settings.

### Western Blots

HEK293T cells (ATCC-CRL-11268) were plated in 12-well plates at 450,000 cells/well density. Cells were transfected 24hrs later via TransIT-2020 (Mirus MIR5400) at 1 µg DNA total/well. 36hrs post-transfection, cells were briefly washed 1x with PBS and 0.5 mL solubilization buffer (20 mM HEPES pH 7.4, 150 mM NaCl, 2 mM MgCl_2_, 0.1 mM EDTA, 2 mM CaCl_2_, 1% Triton X-100) was added to each well. Cells were detached via cell scrapers, transferred to microcentrifuge tubes, and incubated at 4°C with rotation for 30 mins for solubilization. The solubilized cells were centrifuged at 14,000 x g for 15 mins at 4°C and the supernatant was collected as the solubilized cell lysate fraction. Total protein was measured with the BCA assay (Thermo, Pierce #23225). Samples were diluted into sample buffer (312.5 mM Tris-HCl pH 6.8, 10% SDS, 50% glycerol, 12.5% 2-mercaptoethanol, bromophenol blue, and protease inhibitors (Roche Cat# 11873580001), and 3 µg total protein/well was run on 4-20% SDS-PAGE gels (BioRad mini-protean TGX Cat#4561096) at 75 V at room temperature. The Blue Protein Standard, Broad Range (NEB Cat# P7718) was used as a molecular weight ladder. Protein was transferred onto nitrocellulose transfer membrane in transfer buffer (25.1 mM Tris, 192 mM glycine, 20% methanol) at 100 V for 2 hrs at 4°C. Membranes were blocked in 4% bovine serum albumin (BSA, Sigma Cat# 10735086001)/TBST (20 mM Tris-HCl pH 7.4, 150 mM NaCl, 0.05% Tween-20) for 1 hr at room temperature, incubated in primary antibodies diluted into 4% BSA/TBST overnight at 4°C (anti-FLAG M2 mouse, Sigma #F1804, 1:2,000), washed 3 x 5 mins in TBST, incubated in corresponding secondary antibodies (Licor IRDye 800CW donkey anti-mouse Cat#92632212) diluted 1:10,000 into TBST, washed 5 x 5 mins in TBST, and imaged on a Licor Odyssey system. Full length Mus musculus CELSR1 and CELSR2 were used as controls (*70*). Overexpression cDNAs for HEK293T studies were encoded in the pEB Multi-Neo vector (Wako Chemicals, Japan). Cloning and mutagenesis was performed using Gibson assembly and the In-Fusion Assembly system (Takara #638948).

### Surface labeling and immunocytochemistry

HEK293T cells (ATCC #CRL-11268) were maintained in DMEM (Gibco #11995065) with 10% FBS (Gibco #16000044) and 1X penicillin-streptomycin (Corning #MT30002CI) at 37°C and 5% CO2, for a maximum of 25 passages. The FMIL-1 expression construct (pEB HA-FMIL1-FLAG) was generated by cloning HA- and FLAG-tagged *Caenorhabditis elegans* FMIL1 into the pEB multi backbone vector, with the empty vector serving as a control. Primary labeling used rabbit anti-HA (Cell Signaling Technology #3724S; 1:2,000), and secondary labeling used goat anti-rabbit Alexa Fluor 488 (Invitrogen #A-11008; 1:1,000). Glass coverslips in 24-well plates were coated with 100 µL of 50 µg/mL poly-D-lysine (Gibco #A38904-01) for 2 hrs at 37°C. Excess poly-D-lysine was removed, and coverslips were washed with sterile ddH2O. Cells were plated at 1.5 × 10 –2.0 × 10 cells/well in 0.5 mL complete DMEM containing 4-Ara-C (cytarabine; Sigma-Aldrich #C6645; 1:1,000) to inhibit cell proliferation. After 16–24 hrs, cells were transfected with pEB HA-FMIL1-FLAG or the empty pEB multi-vector control via TransIT-2020 (Mirus Bio #MIR5400) with 0.5 µg of total DNA per well in Opti-MEM (Gibco). The transfection mixture was incubated at room temperature for 15–20 min before use, and 50 µL was added dropwise to each well. Cells were incubated at 37°C for 48 hrs before fixation. To distinguish surface-localized from cytoplasmic FMIL-1, cells were fixed and then processed in parallel using one of two labeling protocols: for both protocols, cells were washed once with PBS, fixed in 4% paraformaldehyde (PFA; EMS #15714S)/4% sucrose/PBS for 20 min at 4°C protected from light, and washed 3 × 5 min in PBS; for surface receptor labeling, fixed, non-permeabilized cells were blocked directly in 4% BSA/3% normal goat serum (Fisher #NC9660079)/PBS for 30 min at room temperature, while for total receptor labeling, samples were permeabilized in 0.2% Triton X-100/PBS for 5 min at room temperature and then transferred to blocking buffer (4% BSA/3% normal goat serum (Fisher #NC9660079)/PBS) for 30 min at room temperature. Following blocking, cells from both conditions were incubated with rabbit anti-HA (1:2,000) in blocking buffer for 2 hrs at room temperature, washed 3 × 5 min in PBS, and incubated with goat anti-rabbit Alexa Fluor 488 (1:1,000) together with DAPI (Sigma #10236276001; 1:1,000) and Alexa Fluor 647-conjugated phalloidin (Fisher #50-646-255; 1:40 in methanol) in blocking buffer for 1 hr at room temperature. Cells were washed 3 × 5 min in PBS, and coverslips were mounted on microscope slides using 9 µL ProLong Gold antifade reagent (Invitrogen #P36930) per coverslip.

### Behavioral Analyses

Behavior was assessed using a MBF Biosciences WormLab system. Gravid adults were allowed to lay on NGM plates seeded with 100 µL OP50 containing 1mM all-trans retinal (ATR) in OP50 for 7 hours and then removed. Adult offspring were tested 3 days later. The plates were kept in the dark at all stages. For imaging, the plates were placed on the WormLab imaging platform for 30s before recording started. Video recording began with 10s of baseline recording, followed by 1s on/1s off intervals of red light. Worm movement was processed using Tierpsy Tracker and analyzed using a custom R script (*71*, *72*). Each experiment included 7 plates from each genotype and was repeated on 3 separate days.

### Statistical analysis

Paired comparisons were analyzed using the Student’s t-test. Multiple comparisons were analyzed with a one-way ANOVA with Tukey’s post hoc test using GraphPad Prism 10.

## Results

### Visualizing neuron-specific synapses in the PVD nociceptive circuit with fluorescent markers

A pair of PVD nociceptive neurons arises during the second larval stage (L2) at the postdeirids, bilaterally placed sensory organs positioned on the left and right sublateral nerve cords and posterior to the vulva. PVD morphogenesis initiates with axonal outgrowth into the densely packed ventral nerve cord (VNC) to form glutamatergic synapses with the command interneurons PVC and AVA (*23*). The PVD neurons simultaneously elaborate stereotypically branched dendritic arbors that envelop the body to detect noxious stimuli (Fig 1A, B) (*3*, *32*). With this circuit in place, PVD can trigger an escape response, primarily forward movement, via activation of PVC (*24*).

We used neuron-specific gene expression data from the CeNGEN project to determine that the neuropeptide-encoding gene, *nlp-82*, is selectively expressed in PVC among neurons with processes in the VNC region in which PVD synapses with PVC (Fig 1B) (*61*). A reporter containing the *nlp-82* upstream region, *nlp-82p*::Scarlet::NLG-1 (Neuroligin), produces distinct puncta in the VNC region (posterior to the vulva) in which PVD→PVC synapses are located (*2*). PVD pre-synaptic domains were labeled with GFP::RAB-3 to produce overlapping green (GFP::RAB-3) and magenta (Scarlet::NLG-1) puncta in this region which we identify as PVD→PVC synapses (*73–75*). 24 ± 5 overlapping (green + magenta) puncta are typically detected in L4 animals. 10-20% of the total synaptic puncta are “unpaired” green or magenta puncta, possibly corresponding to other minor synaptic partners for PVD and PVC in this region (Fig 1C, arrowheads) (*3*).

### The MEC-3 LIM-homeodomain transcription factor promotes the formation of PVD synapses with PVC

To initiate a search for genes involved in PVD synapse development, we used Fluorescence-Activated Cell Sorting (FACS) to isolate PVD neurons during the period of PVD axon outgrowth (L2/L3 larval stage) for bulk RNA-sequencing (RNA-seq) (Sup Fig 1A, Sup Table 2) (*33*). We cross-referenced the PVD-enriched transcription factors (TFs) from bulk RNA-seq with the L4 CeNGEN dataset and selected 21 candidates for RNAi screening (see Methods) (Sup Table 3). We observed disruption to the PVD-PVC synaptic marker for only two targets: *lin-39* and *mec-*3. *lin-39* encodes a HOX family transcription factor that promotes cell fate in the midbody region, and PVD fails to elaborate neurites in *lin-39* mutants (*76–78*). Accordingly, we observed no PVD-pre-synaptic GFP::RAB-3 signal and a reduced number of PVC post-synapses in a *lin-39* mutant (data not shown). *mec-3* encodes a LIM class homeodomain protein that functions as a key regulator of PVD fate (*79*, *80*). MEC-3 expression is limited to touch neurons (ALM, PLM, AVM, PVM) and the polymodal nociceptive neurons (FLP, PVD). *mec-3* null mutants largely lack PVD dendritic outgrowth and have a reduced escape response to PVD activation (*24*, *35*). Interestingly, PVD axon outgrowth is not curtailed by *mec-3* loss-of-function (*81*). A *mec-3* null allele showed significantly fewer PVD-PVC synapses than the control (Fig 2A). We quantified the total number of pre-synapses, post-synapses, and noted instances of co-localization (Fig 2B) as well as “unpaired” synapses (Sup Fig 2A). We detected a significant reduction in co-localized, pre-, and post-synapses in *mec-3* mutants. Our results suggest that *mec-3* regulates downstream genes that are necessary for PVD synapses with PVC.

### The MEC-3 target FMIL-1 promotes the formation of PVD synapses with PVC

We next sought to identify genes regulated by *mec-3* for assembly of PVD synapses with PVC. Using FACS, we isolated fluorescently labeled PVD neurons in *mec-3* null mutants for bulk RNA-seq. Comparison to our RNA-seq profile of *mec-3(+)* PVD neurons revealed multiple genes down-regulated in *mec-3* mutant animals (Fig 2C, Sup Table 4). We focused on seven *mec-3*-regulated genes with features suggestive of a potential role in synaptic target selection (e.g. extracellular domains) and combined loss-of-function mutants for each of these genes with our marker for PVD-PVC synapses. The selected targets were: *fmil-1* (GPCR)*, C03A3.1* (unstudied membrane protein)*, kpc-1* (furin homolog)*, seb-2/dhrr-1 (*GPCR*), T03F6.4* (F-box protein), *vab-1* (ephrin receptor)*, and zig-1* (IG family transmembrane protein). Only two of these mutants resulted in a PVD-PVC synaptic defect: *C03A3.1* ablation resulted in a minor increase in PVD→PVC synaptic puncta whereas the deletion allele *fmil-1(gk643828)* caused a significant reduction in PVD→PVC synapses (Fig 2D). We confirmed this result with the deletion allele *fmil-1(ok2115)* which also resulted in dramatically reduced numbers of PVD→PVC synapses (Fig 2E, F, Sup Fig 2B). Together, our findings indicate that the MEC-3 transcription factor promotes expression of *fmil-*1 to establish PVD→PVC synapses.

### FMIL-1 functions cell autonomously in PVD and is enriched in PVD axons

*fmil-1* encodes an adhesion class G protein-coupled receptor (aGPCR), a protein family recently implicated in synaptogenesis (*36*, *37*). At the L4 stage, *fmil-1* transcript is enriched in PVD and in the touch neurons ALM and PVM with lower expression in a few additional neurons (*61*). Transgenic expression of FMIL-1 protein with a C-terminal GFP fusion resulted in strong localization of FMIL-1::GFP signal along the PVD axon with weak expression in PVD dendrites (Fig 3A). In addition, we used CRISPR to tag the endogenous FMIL-1 protein with 7 repeats of the GFP11 fragment and then expressed the GFP1-10 fragment specifically in PVD to reconstitute fluorescent GFP signal (*82*). This split-GFP strategy did not disrupt PVD-PVC synapses, suggesting that the GFP11 tag does not impair FMIL-1 synaptogenic function (Sup Fig 2C). We observed pan-axonal localization of endogenous FMIL-1, confirming our earlier results obtained from transgenic overexpression of FMIL-1::GFP (Fig 3B). In this case, the endogenous FMIL-1 reporter did not result in detectable dendritic FMIL-1::GFP signal, suggesting that FMIL-1 is an axonal protein. Notably, exogenous expression of FMIL-1::GFP protein in PVD fully rescued the PVD→PVC synaptic defect in an *fmil-1* mutant (Fig 3C). While selective expression of FMIL-1::GFP in PVC results in a detectable GFP signal in the PVC region of the ventral nerve cord, PVC expression of FMIL-1::GFP did not rescue the PVD→PVC synaptic defect in an *fmil-1* mutant (Sup Fig 2D,E). Based on these experiments, we propose that FMIL-1 acts pre-synaptically in PVD to promote PVD→PVC synapses.

**Figure 3:**
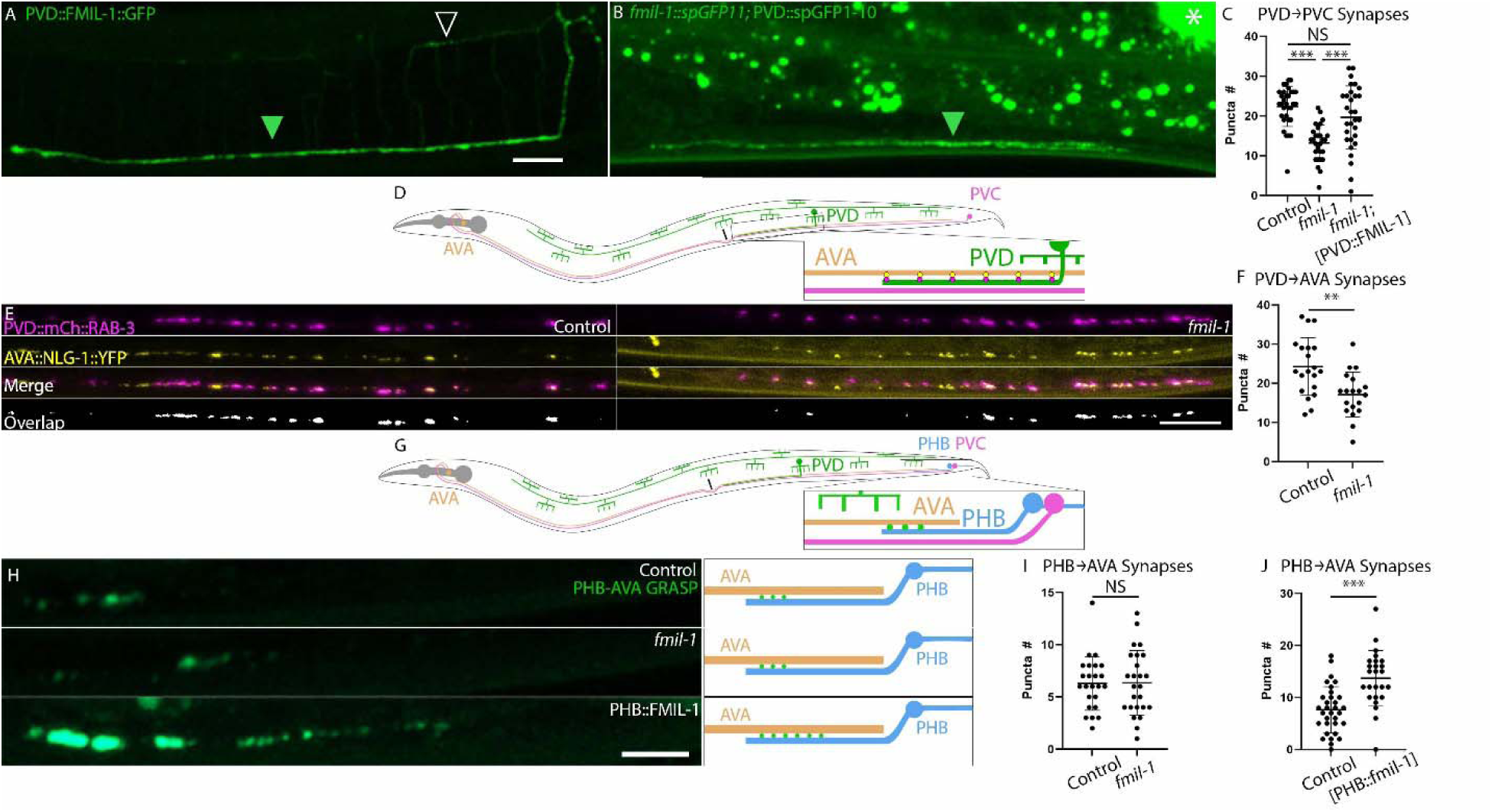
Axonal localization of FMIL-1 promotes synaptogenesis. A. PVD::FMIL-1::GFP is enriched in the PVD axonal compartment (green arrowhead). Dim expression is also visible in PVD dendrites (open arrowhead). Scale bar 10µm. B. Endogenously tagged *fmil-1::gfp11x7* combined with PVD::GFP1-10 shows pan-axonal FMIL-1 expression (green arrowhead). Asterisk denotes coelomocyte::GFP co-injection marker. Punctate gut autofluorescence is visible in region dorsal to PVD axon. Scale bar 10 µm. C. PVD-specific expression of FMIL-1 restores PVD→PVC synapses to *fmil-1* mutant animals. Quantification of PVD→PVC synapses (control v. *fmil-1* p = 2.3x10^-7^, control v. PVD::FMIL-1::GFP, p = 0.19, *fmil-1* v. PVD::FMIL-1::GFP, p = 3.0x10^-4^). n≥28 for all conditions. One-way ANOVA with Tukey’s multiple comparison test. D. Schematic of PVD AVA synaptic labeling strategy between PVD (mCh::RAB-3, magenta) and AVA (NLG-1::YFP, yellow). E. PVD pre-synapses (mCh::RAB-3) and AVA post-synapses (NLG-1::YFP) are reduced in *fmil-1(ok2115)* relative to control, resulting in fewer PVD AVA synapses (Merge and Overlap). Scale bar 10µm. F. PVD AVA synapses are reduced in *fmil-1* nulls (p = 0.0013, n≥19). Student’s t-test. G. Schematic of PHB sensory neuron (blue) with process extending from the PHB soma in the tail to synapses with AVA (tan) that can be visualized using the GRASP reporter (green puncta). H. GRASP labeled PHB AVA synapses are unchanged between control and *fmil-1(ok2115)* but are increased with forced expression of FMIL-1 in PHB. Scale bar 5 µm. Neighboring cartoons (right) depict PHB AVA GRASP signal (green puncta). I. Quantification shows PHB AVA synaptic puncta are unchanged in *fmil-1* nulls (p = 0.84, n=25). Student’s t-test. J. Quantification shows PHB AVA synaptic puncta are significantly increased with forced expression of FMIL-1 in PHB (p = 2.6x10^-6^, n≥24). Student’s t-test.

### PVD and PVC neurons fasciculate normally in fmil-1 mutants

We next considered the possibility that the reduced number of PVD→PVC synapses in *fmil-1* mutants could be due to de-fasciculation between PVD and PVC. To address this question, we labeled PVD with GFP (green) and PVC with Scarlet (magenta) and crossed in the *fmil-1* null mutation (Sup Fig 3A). We categorized fasciculation as either high (no detectable gaps between PVD and PVC axons), medium (occasional short separations between the neurons) or low (extended separations between the neurons). Although the fraction of PVD and PVC axons showing low fasciculation increased from ∼10% in the control to ∼20% in *fmil-1* mutants, the fraction of PVD and PVC axons showing high fasciculation was detectably greater in *fmil-1* mutants (∼50%) than in control animals (∼35%) (Sup Fig 3B). Thus, our results argue against a significant role for FMIL-1 in fasciculation of PVD with PVC and support the hypothesis that FMIL-1 is directly involved in PVD→PVC synaptogenesis.

### FMIL-1 regulates the formation of PVD synapses with AVA

We next considered whether *fmil-1* is also required for PVD synapses with AVA interneurons (Fig 3D). In an EM reconstruction of the ventral nerve cord, PVD makes a similar number of synapses with PVC and AVA and many of these are dyadic for both post-synaptic partners (*2*, *3*). We combined two previously generated extrachromosomal arrays, *PVD::mCherry::rab-3* and *AVA::nlg-1::YFP* to visualize PVD AVA synapses (*8*, *81*). In this experiment, we detected 24 ± 7 dual-colored puncta (mCherry + YFP) in the PVD axonal region posterior to the vulva (Fig 3E, F) in which we have previously observed a comparable number (24 ± 5) of PVD→PVC synapses (Fig 2B). Significantly fewer PVD AVA synapses (17 ± 6, p = 0.0018) were detected in an *fmil-1* null allele indicating that *fmil-1* also regulates PVD-AVA connectivity (Fig 3E, F). This result suggests that FMIL-1 acts in PVD to coordinate synapses with both of its post-synaptic partners, PVC and AVA.

### FMIL-1 expression is sufficient to induce synapse formation in a heterologous circuit

To test the synaptogenic function of FMIL-1, we asked if FMIL-1 is sufficient to induce synapses in other neurons in which FMIL-1 is not normally expressed. We selected PHB AVA synapses to address this question for two reasons. First, PHB AVA synapses are readily detectable with an available GFP Reconstruction Across Synaptic Partners (GRASP) marker (*8*). Second, our results suggest that AVA interacts with FMIL-1 in PVD to create PVD AVA synapses, indicating AVA could also be competent to respond to FMIL-1 expression in other pre-synaptic partners. PHB is a sensory neuron in the tail with an anteriorly directed process that synapses with AVA (Fig 3G). PHB AVA GRASP shows 6 ± 3 synaptic puncta in control conditions. The number of PHB AVA GRASP puncta is unchanged in *fmil-1* mutants (6 ± 3, p = 0.8) (Fig 3H, I), a result consistent with RNA-seq profiles of PHB that failed to detect *fmil-1* transcripts (*61*, *83*). We next used the PHB promoter to drive expression of a *fmil-1::SL2::3xNLS-GFP* and detected significantly elevated PHB AVA GRASP puncta (14 ± 5, p = 2.6x10^-6^) (Fig 3H, J), thus suggesting that FMIL-1 expression in a heterologous pre-synaptic partner (PHB) is sufficient to induce synapses with AVA.

### FMIL-1 acts early in PVD→PVC synapse development

The PVD axon enters the ventral nerve cord soon after the PVD soma first appears in the mid-L2 larval stage (*84*). We used our synaptic marker to image PVD→PVC dual-colored synaptic puncta in early L3 larvae to study how PVD→PVC synapses are affected by FMIL-1 at the onset of axonal growth (Fig 4A). We reasoned that If FMIL-1 promotes synapse formation, then fewer PVD→PVC synapses should be detected in an *fmil-1* mutant at this early larval stage but should remain steady over time. In contrast, if FMIL-1 functions strictly to maintain synapses after they are established, the number of PVD→PVC synapses in an *fmil-1* mutant should be initially equivalent to control but then decrease during later development.

**Figure 4:**
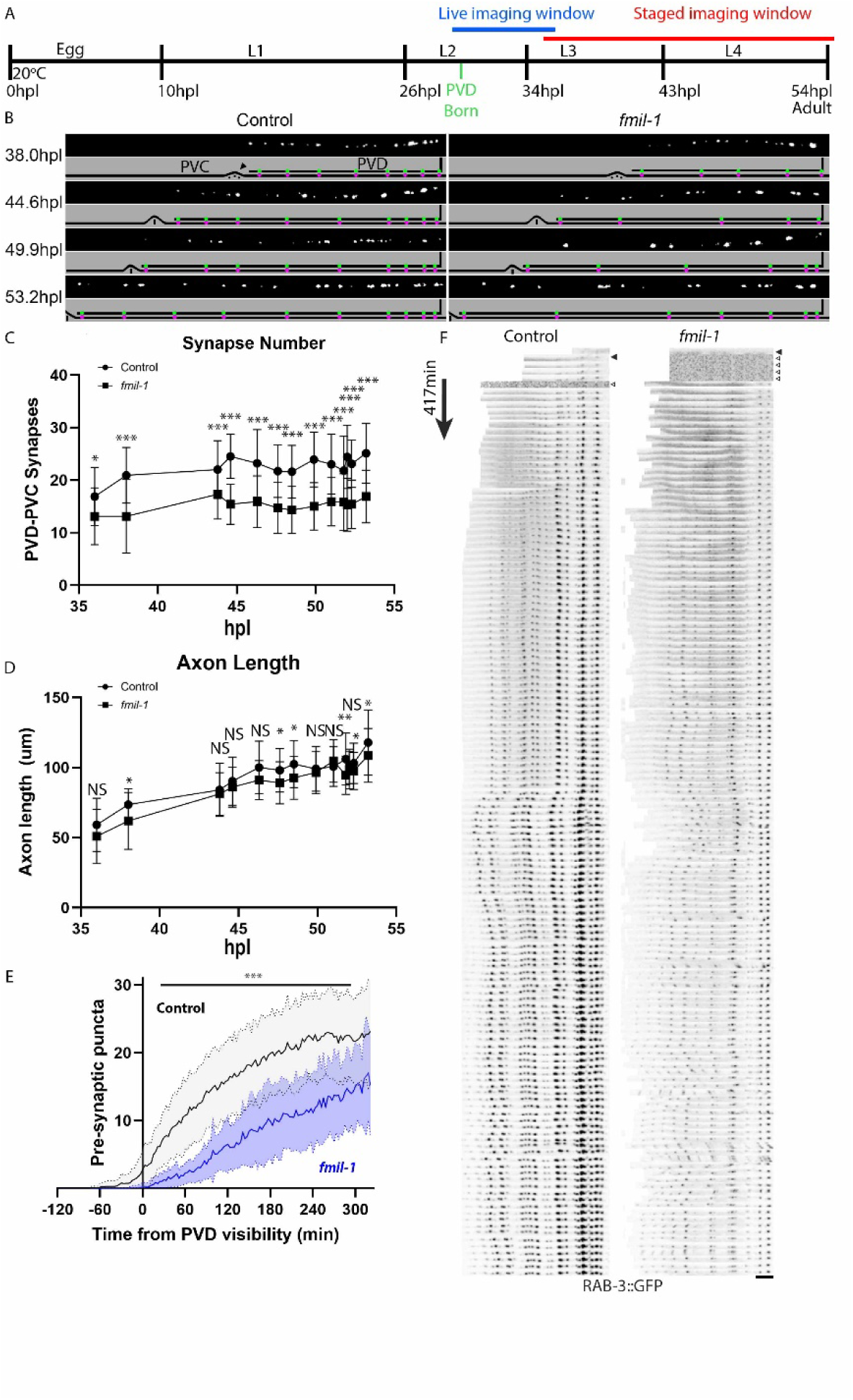
Time-lapse imaging of PVD→PVC synaptogenesis reveals a role for *fmil-1* in synapse formation vs maintenance. A. *C. elegans* developmental timeline. 0 hours post-lay (hpl) marks the time of egg laying. Note PVD birth in the L2 stage (green), live-imaging window of mid L2-mid L3 (blue) and imaging window of early L3-adult (red). B. Representative time points for PVD→PVC synapses in control (left) and *fmil-1* (right). (Top) Overlapping GFP::RAB-3 (PVD) and Scarlet::NLG-1 (PVC) puncta (white). (Bottom) Schematic of PVD axon with GFP::RAB-3 puncta (green) vs PVC axon with Scarlet::NLG-1 puncta (magenta). The PVD axon reaches its anterior limit at the vulva in L3 (38 hpl, arrowhead) but continues to lengthen as the larval animal increases in size. C. PVD→PVC synapse number during larval development for control vs *fmil-1(ok2115)*. Error bars mark SD. Statistical comparisons are control vs *fmil-1* at each time point, * p = 0.03, ***p < 0.001, Student’s t-test. See Supplemental Table 5 for all statistical results. D. Growth of PVD axon during larval development for control vs *fmil-1(ok2115)*. Error bars mark SD. *p < 0.05, **p < 0.01, NS = Not Significant, Student’s t-test. See Supplemental Table 5 for all statistical results. E. Quantification of PVD::GFP::RAB-3 in control (black) and *fmil-1* (blue) over 6 hours. PVD::myr::mCherry is first visible in the PVD cell body at time 0. Timepoints are n≥10 for both conditions. ***p < 0.001 for control vs *fmil-1* at all time points under horizontal black bar by Student’s t-test. See Supplemental Table 5 for all statistical results. F. Representative time series of PVD::GFP::RAB-3 accumulation in control (left) and *fmil-1* (right). Black arrowhead denotes first frame in which the PVD cell body (not pictured) is visible. White arrowheads denote frames with out-of-focus GFP::RAB-3 puncta. Imaging frequency is three minutes and montage covers 417 minutes of development. Scale bar 10 µm.

To distinguish between these alternative roles for FMIL-1 of synapse formation versus maintenance, we quantified PVD→PVC synapses in the L3 stage when the PVD axon enters the ventral nerve cord. These measurements detected significantly fewer PVD→PVC synapses in the *fmil-1* mutant vs control (Fig 4B,C). To determine if synapse number changed over time, we compiled PVD→PVC synaptic counts for wild-type and *fmil-1* animals, using vulval morphology to order developmental age (*47*). PVD→PVC synaptic counts were consistently lower for *fmil-1* mutants than control throughout the L4 larval period. Even the earliest timepoint of 36hpl showed *fmil-1* mutants with fewer puncta (13 ± 5) than control animals (17 ± 5, p = 0.03). In addition, synaptic number appeared largely stable throughout the L4 period of PVD axonal outgrowth, with no significant increase in synapse number from mid-L3 (38hpl) to late L4 stages (52.3hpl, wt p = 0.07, *fmil-1* p = 0.07) (Fig 4C). By contrast, PVD axonal length increases by ∼2-fold for both *fmil-1* (p = 8.7x10^-15^) and control (p = 2.4x10^-13^) animals during this period (Fig 4D). Our results are consistent with the hypothesis that FMIL-1 promotes the formation of PVD→PVC synapses but is likely not required for their maintenance. In addition, we determined that total PVD axon length increased to similar extents during the L3-L4 larval stages for both *fmil-1* and control animals (Fig 4D) which suggests that *fmil-1* is not required for PVD axonal outgrowth.

As an additional test of the hypothesis that FMIL-1 acts at the earliest stages of synaptogenesis, we used time-lapse imaging to observe the formation of PVD→PVC synapses. In this experiment, the PVD axon, labeled with *ser2prom3::myr::*mCherry, was initially visible immediately before the transition from the L2 to L3 larval stages. *ser2prom3::*GFP::RAB-3 puncta are first visible at approximately the same time as the PVD cell body. Movies were collected over 8 hours in both control and *fmil-1(ok2115)* mutants (Sup Movies 1-4). We quantified the GFP::RAB-3 puncta at all time points and confirmed that synapse formation is more rapid and plateaus at a higher level in controls than in *fmil-1* mutants (Fig 4E). The earlier appearance and brighter signal for GFP::RAB-3 puncta in controls vs *fmil-1* animals is strikingly evident in a montage of these movies (Fig 4F). Overall, these findings support the hypothesis that FMIL-1 acts during PVD axon outgrowth to promote synapse formation.

### fmil-1 encodes an adhesion G protein-coupled receptor (aGPCR)

Based on sequence homology and AlphaFold modeling, the FMIL-1 protein is a member of the adhesion GPCR family. aGPCRs are defined by a large extracellular region that includes an autoproteolysis-inducing (GAIN) domain (*85*). Although GAIN domains are typically cleaved in the ER, the N-terminal fragment (NTF) remains non-covalently attached to the C-terminal fragment (CTF) as they traffic together to the plasma membrane. Either adhesive binding or shearing force is thought to separate the NTF from the CTF and thereby activate aGPCR signaling via the newly exposed tethered agonist (TA). Alternatively, the intact NTF/CTF complex may also activate aGPCR signaling (*86*, *87*). In some cases, the NTF fragment can function as an independent juxtacrine signal to adjacent cells (*88*, *89*). The FMIL-1 extracellular domain (ECD) contains cadherin repeats (Cad1, Cad2), a Fibronectin-type III domain (Fn), an EGF domain and a hormone receptor-like domain (HormR). These putative binding domains are linked to the GAIN domain and the 7-pass transmembrane domain. The FMIL-1 intracellular domain of 100aa has no known homology and is predicted to adopt an unstructured conformation (Fig 5A). FMIL-1 is most similar to the CELSR family of aGPCRs, due to the combination of cadherin, EGF and hormone receptor domains (*90*). Notably, FMIL-1 (1,029aa) is considerably smaller than other members of the CELSR family including FMI-1 (2,596aa) in *C. elegans*, Starry Night (*stan*, 3,579aa) in *Drosophila* and CELSR1, CELSR2, and CELSR3 in humans (3,014aa, 2,923aa, and 3,312aa respectively).

**Figure 5:**
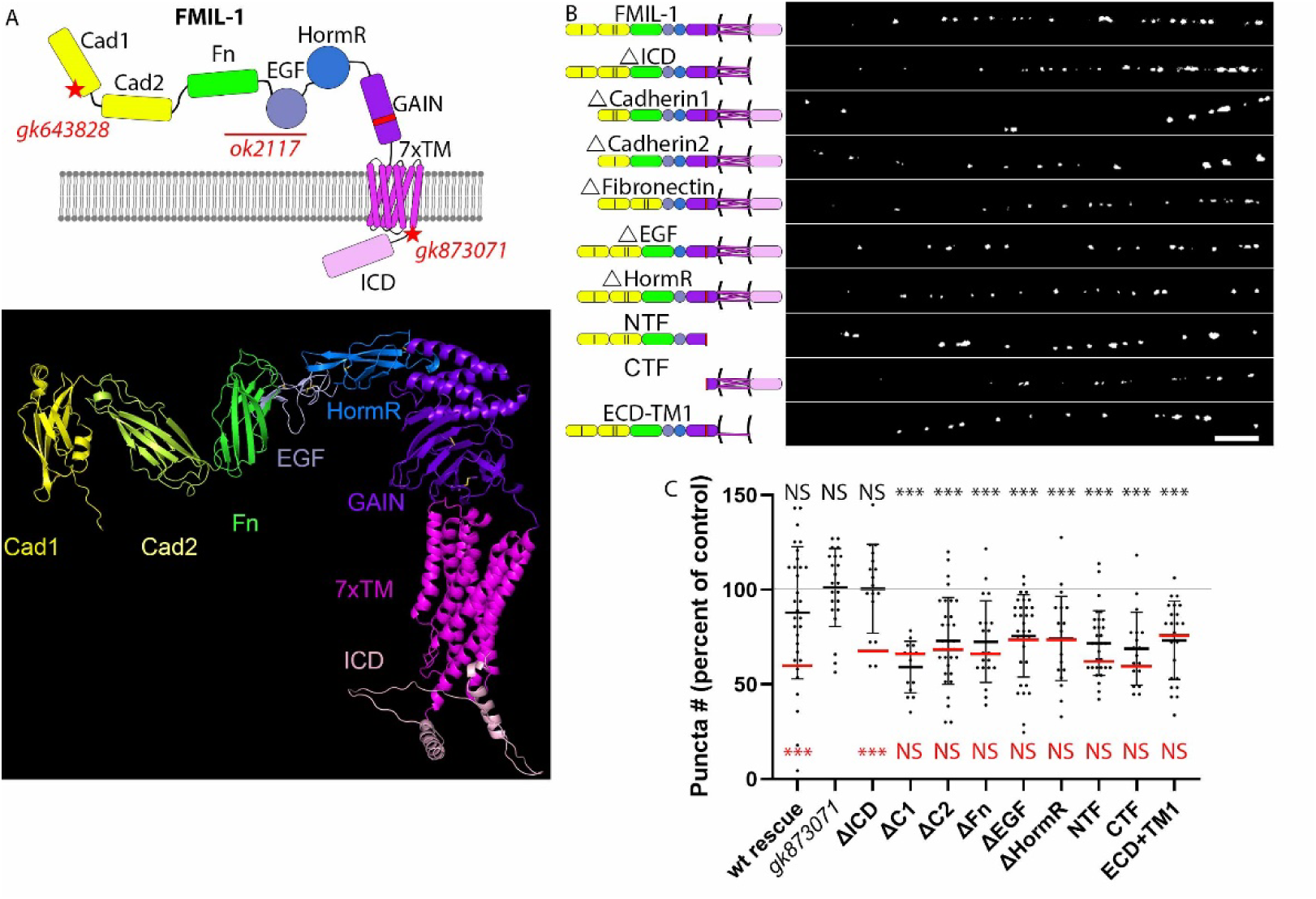
FMIL-1 Extracellular Domain (ECD) but not Intracellular Domain (ICD) is required for PVD→PVC synaptogenesis. A. (Top) FMIL-1 is an adhesion-class GPCR with the following domains: cadherin (Cad1, Cad2) (yellow), fibronectin type-III (green), EGF (gray), hormone receptor (blue), GAIN (purple, red bar denotes cleavage site), 7-pass transmembrane (pink), and intracellular (light pink). Genomic mutations: *gk643828* (splice-donor), *ok2115* (deletion frameshift), *gk873071* (premature stop in final exon). (Bottom) AlphaFold structure for FMIL-1. B. (Left) FMIL-1 domains and mutant constructs tested for rescue of PVD→PVC synaptic defect in *fmil-1(ok2115)*. (Right) Overlapping of PVD::GFP::RAB-3 and PVC::Scarlet::NLG-1 puncta (white) for each FMIL-1 construct. Scale bar = 10 µm. C. Quantification of PVD→PVC synapses. Black horizontal lines denote mean puncta number ± SD for each experiment and red lines denote mean puncta number for *fmil-1(ok2115)* mutant paired with each *fmil-1* mutant construct. Black lettering at top shows statistical results for wild-type control v. FMIL-1 rescue constructs and genomic mutants; Red lettering at bottom shows statistical results for *fmil-1(ok2115)* vs. FMIL-1 rescue constructs. ***p < 0.001, NS = Not Significant, one-way ANOVA with Tukey’s post-hoc multiple comparison test. Data for the FMIL-1 wild-type rescue construct are replotted from Figure 3C. See Supplemental Table 5 for all statistical results.

### The FMIL-1 intracellular domain (ICD) is not required for PVD→PVC synapse formation

To test for a functional role for the FMIL-1 intracellular domain (ICD), we used a mutant *fmil-1(gk873071)* with a premature stop codon that truncates the ICD to 6 amino acids proximal to the transmembrane domain (*91*). *fmil-1(gk873071)* did not show a PVD→PVC synaptic phenotype and was indistinguishable from controls (Fig 5C). To confirm this result, we fused GFP with a flexible linker to the C-terminus of the truncated *fmil-1(gk873071)* protein (FMIL-1::ΔICD::GFP) and expressed it in the *fmil-1* null background. FMIL-1::ΔICD::GFP trafficked to the axon, suggesting proper folding and expression (Sup Fig 4A), and fully rescued the PVD→PVC synaptic defect (Fig 5B, C). These results suggest that the FMIL-1 ICD region is not required for FMIL-1-dependent synaptogenesis, a finding that differs from previous results showing that the aGPCR ICD region is required for downstream signal transduction (*92*, *93*).

### FMIL-1 extracellular domains are required for synaptogenic function

To assess the *in vivo* function of the FMIL-1 ExtraCellular Domain (ECD), we removed each of the identified ECD subdomains (Cad1, Cad2, Fn, EGF, HormR) to produce a series of FMIL-1 structural variants (Fig 5B). Each construct was fused to GFP at the C-terminus and transgenically expressed in PVD in an *fmil-1(ok2115)* null mutant. We confirmed that each of these constructs localized to the PVD axon (Sup Fig 4A). Interestingly, none of these FMIL-1 ECD variants rescued the *fmil-1* mutant PVD→PVC synaptic phenotype (Fig 5C). These results suggest that the entire FMIL-1 ECD is required for its synaptogenic function.

To assess the activity of a potentially cleaved form of FMIL-1, we fused Scarlet to an N-terminal fragment mimicking the predicted NTF and expressed it in PVD. The truncated FMIL-1-NTF::Scarlet protein failed to restore PVD→PVC synapses to an *fmil-1* mutant, a result arguing against its function as a juxtacrine signal in PVD (Fig 5B, Sup Fig 4A). We next tested a pre-cleaved CTF to determine if it could function as a presumptive tethered agonist (TA) for FMIL-1. Transgenic expression of the FMIL-1-CTF failed to restore PVD→PVC synapses to an *fmil-1* mutant. This result suggests that the canonical TA model of aGPCR function is not sufficient for PVD→PVC synaptogenesis. Finally, we tested the *in vivo* activity of the intact FMIL-1 ECD region. A construct containing the FMIL-1 ECD and first transmembrane domain was transgenically expressed in PVD, but it also failed to rescue the fmil-1 synaptic defect (Fig 5B, C).

Although each of the FMIL-1 ECD constructs is likely overexpressed, all fail to complement the *fmil-1* synaptic defect, which suggests that these mutant forms do not retain residual function. Our findings indicate that FMIL-1 function depends on all five of its extracellular adhesion domains. In addition, the NTF seems unlikely to act as a juxtacrine signal and its dissociation at the GAIN cleavage site is not sufficient to trigger synaptogenesis. Finally, our results suggest that the structure of the FMIL-1 transmembrane domain is necessary for function, though the ICD is dispensable, a finding consistent with the observation that the G protein binding domains in GPCRs are often embedded in the 7-TM intracellular loops (*94*).

### FMIL-1 synaptogenesis does not rely on cleavage by the GAIN domain

As noted above, the “pre-cleaved” NTF and CTF fragments do not rescue the *fmil-1* synaptic phenotype, suggesting that the dissociated FMIL-1 protein is inactive. The GAIN domain of cleaved aGPCRs typically contains an H/RXS/T motif at the cleavage site. Histidine or another proton acceptor triggers cleavage through deprotonation of the threonine/serine residue, which then performs a nucleophilic attack to cleave the peptide bond between X and S/T residues (*95–97*). In contrast, aGPCRs that are not cleaved (e.g., CELSR1 and CELSR3) substitute either an alanine or glycine for the canonical serine/threonine residues at the cleavage site. FMIL-1 includes a glycine residue at this position, suggesting that it is not cleaved (Fig 6A) (*70*). To test this possibility, we expressed FMIL-1 and mouse CELSR1 and CELSR2 proteins tagged with N-terminal HA and C-terminal FLAG epitopes in human HEK293T cells. After verifying FMIL-1 trafficked to the cell surface (Sup Fig 5A), we immunoblotted the HEK293T cell extracts and detected full-length proteins for all three constructs, FMIL-1, CELSR1 and CELSR2. In addition, we detected a CTF for CELSR2 which retains the HXS/T motif and is known to be cleaved. In contrast, CELSR1, which substitutes an alanine at the S/T cleavage site does not produce a CTF as expected from previous results (Fig 6B) (*70*). Although FMIL-1 produced a cleavage product of the predicted size, most of the wild-type FMIL-1 protein was intact (Fig 6B). We wondered if T655, a threonine residue adjacent to the conserved histidine (H657), could be required for FMIL-1 cleavage. To test this idea, we generated FMIL-1 mutants in which the threonine residue was converted either to glycine (T655G) or alanine (T655A). Similar mutations (H657G and H657A) were introduced for histidine. A cleavage product was detected for T655A but not for the other FMIL-1 mutants (T655G, H657G, H657A). These results suggest that some proportion of the wild type FMIL-1 protein is cleaved and that this process is mediated by residues T655 and H657. To determine if GAIN domain cleavage is required for FMIL-1 function *in vivo*, we used CRISPR to generate T655G and H657G point mutations in the endogenous *fmil-1* locus. Neither the T655G nor H657G mutations disrupted the formation of PVD→PVC synapses (Fig 6C-D). Thus, our findings suggest that FMIL-1 cleavage is possible, but not necessary for its function, a property shared with other aGPCR family members which have been shown to be either cleavage deficient or cleavage independent (*70*, *87*, *98*, *99*).

**Figure 6:**
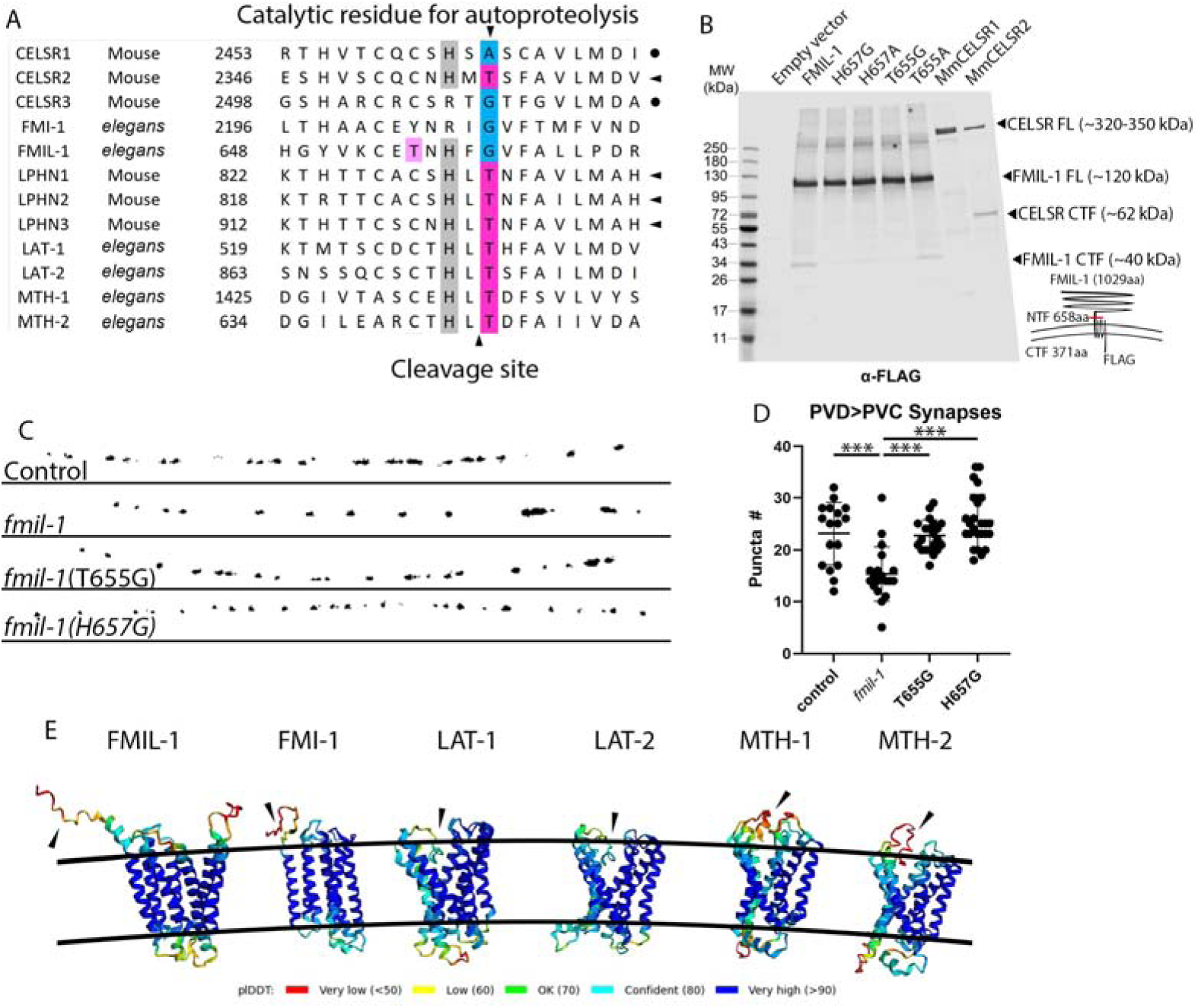
Cleavage at the GAIN domain is not required for FMIL-1 synaptogenic function. A. Alignment of mouse and *C. elegans* aGPCRs in the GAIN cleavage region. Top arrowhead marks the catalytic residue for autoproteolysis and bottom arrowhead indicates cleavage site. Side arrowheads denote known cleaved aGPCRs and circles mark cleavage-deficient proteins. Numbers indicate position of first amino acid. Dark pink highlights conserved threonines that catalyze cleavage. Light pink highlights potential catalytic threonine in FMIL-1. Blue highlights sites where threonine is not conserved. B. Immunoblot of HEK293T cells expressing an empty vector or C-terminal FLAG-tagged aGPCR constructs. Cartoon depicts amino acid (aa) length of intact FMIL-1 vs CTF and NTF fragments. Estimated sizes are shown for each protein. C. Overlay of PVD::GFP::RAB-3 and PVC::Scarlet::NLG-1 puncta shows PVD→PVC connectivity in control, *fmil-1(ok2115)*, *fmil-1(T655G) and fmil-1(H657G)*. D. Quantification of PVD→PVC synaptic puncta. Control v. *fmil-1* p = 6.2x10^-5^, *fmil-1* v. T655G p = 7x10^-5^, *fmil-1* v. H657G p = 1.2x10^-8^ all other comparisons were NS. n≥17 for all conditions. One-way ANOVA with a Tukey’s post-hoc multiple comparison test. See Supplemental Table 5 for all statistical results. anE. AlphaFold prediction of cleaved *C. elegans* aGPCRs. Black arrowheads denote the tethered agonist (TA). Note that the TA is unbound in FMIL-1 and FMI-1 whereas the TA is buried in the TM region in LAT-1, LAT-2, MTH-1 and MTH-2. The ICD has been excluded for simplicity.

aGPCR cleavage is proposed to reveal a Tethered Agonist (TA), which can activate downstream G protein-coupled signaling and arrestin recruitment (*93*, *99*). Structural modeling with AlphaFold has confirmed that the TA for cleavage-dependent aGPCRs is buried in the transmembrane domain whereas the TA for cleavage-independent GPCRs is unbound (*99*). Among AlphaFold models of *C. elegans* aGPCRs, LAT-1, LAT-2, MTH-1 and MTH-2 exhibited a buried TA whereas FMIL-1 and FMI-1 did not show a buried TA in the TM region (Fig 6E). Interestingly, FMIL-1 and FMI-1 include a glycine at the cleavage site whereas all the other C. elegans aGPCRs include the threonine residue characteristic of cleaved aGPCRs. Thus, a structural model of FMIL-1 is also consistent with the hypothesis that FMIL-1 autocatalytic cleavage is not required for its synaptogenic function.

### Predicted binding partners of the FMIL-1 extracellular domain are not required for its synaptogenic function

To identify potential interacting partners for the FMIL-1 ECD, we first considered proteins recently reported to bind the FMIL-1 ECD in an *in vitro* adhesion assay (*100*). Candidates from this screen include the transmembrane proteins PTRG-1 (protein tyrosine phosphatase), SOL-1 (glutamate receptor accessory protein), UNC-40 (DCC, netrin receptor), and the diffusible ligands GPLA-1 (glycoprotein hormone), NDNF-1 (neuron-derived neurotrophic factor ortholog), LECT-2 (chemotaxin homolog), and SCL-10 (GLIPR1 homolog). Loss-of-function mutants for these FMIL-1 interacting proteins failed to cause PVD→PVC synaptic defects with two exceptions, *unc-40,* which showed a strong loss of PVD→PVC synapses (see below), and *sol-1,* which displayed a weak effect that we did not investigate further (Sup Fig 6A).

Genetic ablation of the *unc-40* receptor or its canonical ligand *unc-6* (netrin) reduced PVD→PVC connectivity to approximately the same extent as *fmil-1* null alleles whereas *unc-5* (netrin receptor) loss had no effect (Sup Fig 6A) (*101*). Given the well-established roles of UNC-6/netrin and UNC-40/DCC in axonal guidance, we considered the possibility that the reduced number of PVD→PVC synapses in *unc-6* and *unc-40* mutants could result from defasciculation of the PVD and PVC axonal processes. To investigate this possibility, we labeled PVD with GFP and PVC with Scarlet for 2-color imaging of their axonal processes. In the wild-type, PVD and PVC processes are closely apposed in the ventral nerve cord (Sup Fig 6B-D). In contrast, in *unc-40* and *unc-6* mutants, both PVD and PVC showed numerous pathfinding errors including truncated PVD axons with no PVC fasciculation, missing PVD or PVC axons (Sup Fig 6C), or PVC axons misrouted into lateral nerve cords (Sup Fig 6D). Given the severity of these axonal guidance defects, it was not possible to reliably confirm a separate role for UNC-40/DCC in PVD→PVC synaptogenesis vs pathfinding guidance. We also examined a mutant allele of the protein tyrosine phosphatase *clr-1,* which promotes PHB AVA synapses in conjunction with UNC-40 and UNC-6 (*102*). The *clr-1* mutant did not prevent induction of FMIL-1-dependent PVD→PVC synapses (Sup Fig 6A). Overall, we determined that UNC-40 and UNC-6 guide PVD and PVC pathfinding but could not fully rule out a role for UNC-40 and UNC-6 in interactions with FMIL-1 to promote synaptogenesis.

We next considered possible binding partners for the FMIL-1 cadherin domains, which promote the formation of PVD→PVC synapses (Fig 5C). Our results showing that FMIL-1 expression in PVD is sufficient for PVD→PVC synaptogenesis suggest that homophilic FMIL-1 cadherin interactions are not necessary for PVD→PVC synapse induction. As an alternative model in which heterophilic binding with FMIL-1 cadherin domains promotes PVD→PVC synapse formation, we considered cadherin family proteins *cdh-4*, *casy-1* and *fmi-1* that are known to be expressed in PVC. However, null alleles of *cdh-4*, *casy-1,* and *fmi-1* did not disrupt PVD→PVC synapses. In *fmi-1* nulls, we observed frequently misrouted PVD axons as previously reported (Sup Fig 6E) (*103*). To identify potential binding partners for the FMIL-1 fibronectin domain, we tested viable hypomorphs for the alpha subunit *ina-1* and beta subunit *pat-3,* but neither resulted in a PVD→PVC synaptic defect (Sup Fig 6A). Other integrin subunits are either not expressed in neurons (*inb-1*) or do not have viable mutant alleles (*pat-2*). Finally, we considered Frizzled receptors that function in the planar cell polarity (PCP) pathway, which is known to interact with CELSRs in mammals. We tested loss-of-function alleles for frizzled receptors expressed in PVC, *lin-17, cfz-2,* and *vang-1*, and, but did not observe PVD→PVC synaptic defects (Sup Fig 6A) (*78*). Although our genetic tests did not reveal a candidate adhesion protein for mediating transcellular interaction of FMIL-1 in PVD with the PVC post-synaptic membrane, this model seems very likely, given our robust findings that FMIL-1 is cell autonomous for PVD and that all the cognate protein-protein interaction motifs in the FMIL-1 ECD are required for its synaptogenic function.

### FMIL-1 is necessary for sustained behavioral response to PVD activation

Previous work determined that optogenetic stimulation of PVD triggers forward movement via PVC (*24*). We wondered if loss of PVD→PVC synapses impaired this behavior. In our experiments, red light excitation of PVD::Chrimson resulted in sharp forward acceleration (Fig 7A) (*104*). The PVD→PVC circuit functions as an escape response in which nociceptive input from PVD relayed to PVC evokes rapid movement away from the stimulus (*24*). Although the strength (*i.e.* forward speed) of the response declined with repetitive stimulation (1 sec on + 1 sec off), control animals consistently showed a stronger response than either of the *fmil-1* null alleles tested in this paradigm (Fig 7B). Our results suggest that the reduction in PVD→PVC synapse number in *fmil-1* mutants attenuates the response to sustained stimulation.

**Figure 7:**
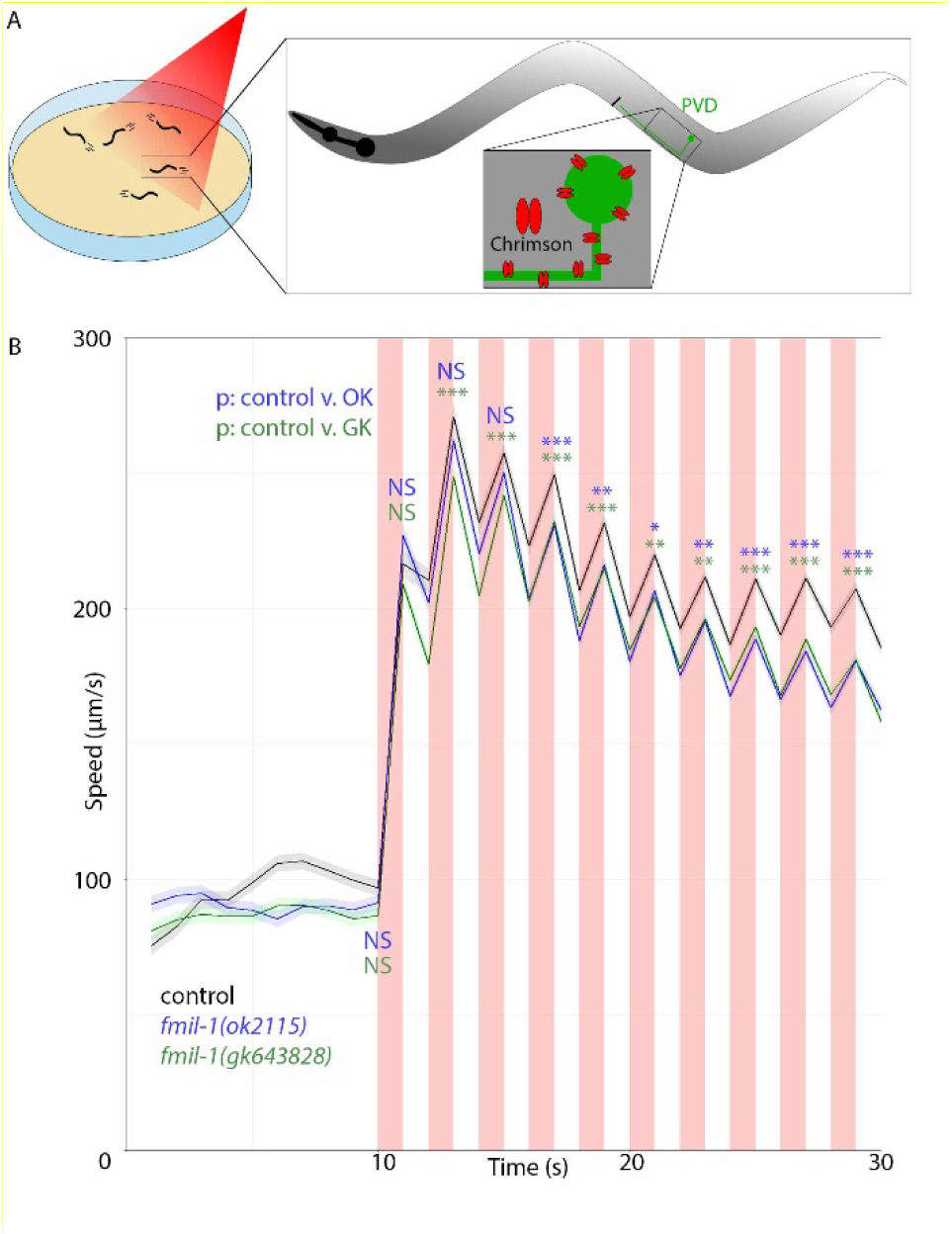
FMIL-1 is required for a robust escape response triggered by optogenetic activation of PVD nociceptive neurons. A. Diagram of optogenetic design to detect PVD nociceptive function. Animals on a culture plate respond to red light by accelerating. Inset: diagram of PVD with axon showing pan-expression of Chrimson. B. Average locomotory speed (µm/s) over time for PVD::Chrimson control (black), *fmil-1(ok2115)* (blue), and *fmil-1(gk643828)* (green) animals. Lighter colors denote SEM for each sample. Vertical red bars denote period (1 s) of red light stimulation. See Supplemental Table 5 for all statistical results.

## Discussion

The circuit architecture of the brain is defined by genetically encoded pathways that direct the creation of neuron-specific synapses. Here we exploit the genetic tractability and ready access to live cell imaging of *C. elegans* to identify a transcriptionally regulated adhesion protein that controls synapse formation in a nociceptive circuit. We show that the LIM-homeodomain protein, MEC-3, and its downstream effector, FMIL-1, a member of the aGPCR family, promote the formation of synapses between the nociceptive neuron PVD and its post-synaptic partners, PVC and AVA (Fig 8). The absence of these FMIL-1-dependent synapses impairs the escape response triggered by nociceptive signals from PVD. The role of the FMIL-1 protein in synaptogenesis is cell autonomous for PVD where it localizes to the axonal membrane. The potent synaptogenic function of FMIL-1 is underscored by our finding that ectopic expression of FMIL-1 is sufficient to induce synapse formation in another sensory circuit. The extracellular adhesion domains of FMIL-1 are characteristic of the CELSR family of aGPCRs, and each is required for FMIL-1 function in PVD→PVC synaptogenesis. In contrast, the FMIL-1 autocleavage activity and its intracellular domain (ICD) are dispensable. Our results highlight the PVD nociceptive circuit as an accessible system for investigating synaptic partner selection and as a new model for understanding how aGPCRs function to direct synaptogenesis.

**Figure 8.**
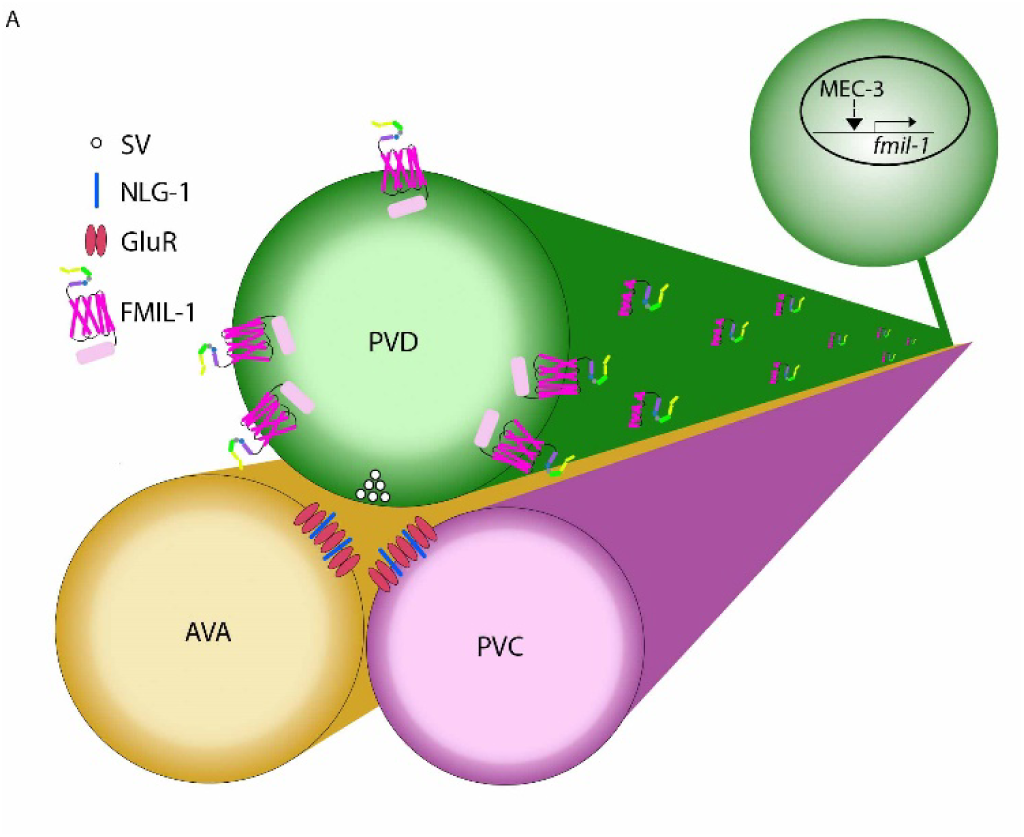
Model of FMIL-1 expression and synaptogenic function in PVD nociceptive neurons. A. The LIM-homeodomain transcription factor MEC-3 activates fmil-1 expression. FMIL-1 protein localizes to the PVD axon in the ventral nerve cord to promote formation of pre-synapses in PVD, assessed here by GFP::RAB-3 localization to synaptic vesicles (SVs), and post-synapses in AVA and PVC command interneurons measured by neuroligin (NLG-1). Although not measured here, glutamate receptors (GluR) cluster along PVC and AVA and likely appose the glutamatergic synaptic vesicles in PVD.

### The FMIL-1 ECD functions pre-synaptically in PVD to promote post-synaptic connections with PVC

aGPCRs have emerged as important molecules in neuronal partner selection and are conserved across species (*38*, *105*). In *C. elegans*, FMI-1/Flamingo, a paralog of FMIL-1/Flamingo-like, has been shown to regulate axon guidance (*106*, *107*) and synapse formation (*108*). FMI-1 and FMIL-1 are members of the CELSR sub-family of aGPCRs that promote synaptogenesis in mammalian hippocampal neurons similar to other aGPCRs, including the latrophilins (*37*). The FMIL-1 ECD contains six distinct domains (2 cadherin, 1 fibronectin, 1 EGF, 1 hormone receptor and 1 GAIN). Removal of any single ECD domain abrogated FMIL-1-dependent formation of PVD→PVC synapses, suggesting that each of these ECD regions is necessary for function. In contrast, subsets of extracellular domains in FMI-1/Flamingo and LAT-1 are not required in vivo (*98*, *106*). Our results could mean that each FMIL-1 extracellular domain binds to a unique ligand and that every interaction is necessary for FMIL-1 function in *C. elegans*. Alternatively, selected domains may be required for ligand binding with the remaining regions, providing structural support.

We considered a heterotypic binding model, as reported for vertebrate aGPCR Lphn3/ADGRL3 interaction with teneurins and FLRTs (*105*). We tested mutants for proteins that interact with the FMIL-1 ECD in an *in vitro* binding assay (SOL-1, PTRG-1, UNC-40, GPLA-1, NDNF-1, LECT-2, SCL-10) (*100*). We also considered cadherins with known expression in PVC (CDH-4, CASY-1, FMI-1), as well as integrins and PCP components (*109*). Null mutants for each of these candidates did not significantly disrupt PVD→PVC synaptic connectivity, with the exception of *unc-40/DCC* which caused PVD and PVC pathfinding defects (see below). Overlapping roles for FMIL-1 binding partners may account for the absence of clear mutant effects (*56*). This possibility could be addressed by the simultaneous removal of multiple targets in future studies. Although UNC-40 directly promotes synaptogenesis in other *C. elegans* circuits (*8*, *110–112*), a potentially separate role for UNC-40 in the formation of PVD→PVC was masked by the misplacement of PVD and PVC axons in *unc-40* mutants. Although homophilic interactions have been reported for FMI-1 and other cadherins (*103*, *113*), this mechanism seems unlikely for FMIL-1 for two reasons. First, *fmil-1* expression in PVC has not been detected in RNA-seq data from multiple developmental timepoints including the embryo, L1, L4 larvae and young adult (*61*, *83*, *114–116*). Second, PVD-specific expression of FMIL-1 is sufficient to rescue the *fmil-1* PVD→PVC synaptic defect (Fig 3C). Taken together, our results suggest that FMIL-1 interacts with a yet unknown heterophilic partner in PVC and AVA to trigger synapse formation.

### Autocatalytic cleavage of the extracellular domain is not required for FMIL-1 synaptogenic function

The GAIN domain, a unique structural region of the ECD, facilitates aGPCR self-cleavage in the endoplasmic reticulum (ER). The resulting N-terminal and C-terminal fragments (NTF/CTF) are trafficked to the cell membrane as a noncovalently bound complex. Extracellular binding is proposed to reveal the N-terminus of the CTF, or tethered agonist (TA), allowing interaction with the transmembrane domain region to activate downstream aGPCR signaling (*87*). However, some aGPCRs are cleavage deficient or do not require cleavage for activation and function (*70*, *98*).

Although the FMIL-1 GAIN domain does not include a canonical autocatalytic cleavage site, expression in mammalian HEK293T cells resulted in partial cleavage that was prevented by mutations that altered a nearby histidine or threonine residue. Neither mutant, however, showed a PVD→PVC wiring defect *in vivo*, suggesting that cleavage is not required for FMIL-1 synaptogenic function. In addition, structural modeling revealed that the TA component of the predicted FMIL-1 CTF fails to bind the transmembrane region as observed for other TA-dependent aGPCRs. Because the ECD is required for FMIL-1 function, we suggest that FMIL-1 may act as a conventional GPCR with an extracellular ligand that triggers a conformational change for activation of internal G protein signaling pathways.

### FMIL-1 promotes synapse formation but is not required for synaptic maintenance

Although a wide array of cell surface adhesion proteins have been proposed to drive synapse formation, *in vivo* confirmation has been rarely achieved largely due to the challenge of distinguishing between synapse induction vs maintenance (*117*). To address this question for FMIL-1, we used live cell imaging to monitor PVD→PVC synapses during PVD axonal outgrowth. We determined that the *fmil-1* synaptic defect occurs soon after the PVD neuron is generated during early larval development and as the PVD axon extends in the ventral nerve cord. Fewer PVD→PVC synapses were observed in an *fmil-1* mutant vs the control at the earliest observable stage. Furthermore, we did not detect any additional loss of PVD→PVC synapses throughout larval development in *fmil-1* mutants, a finding indicating that FMIL-1 is not required for synaptic maintenance. The proposed synaptogenic function for FMIL-1 is supported by our finding that ectopic FMIL-1 expression in the PHB sensory neuron is sufficient to enhance synapse formation with AVA in the ventral nerve cord. We note that the rapid formation of PVD→PVC synapses during PVD axonal outgrowth mimics that of PDE, a dopaminergic neuron that arises from a common lineage with PVD and also simultaneously extends its axon into the ventral nerve cord during early larval development (*118*).

### FMIL-1 functions in concert with additional connectivity determinants for PVD→PVC synaptogenesis

Although MEC-3 and its downstream effector FMIL-1 promote the formation of PVD→PVC synapses, approximately 40% of PVD→PVC synapses are generated in the absence of either *mec-3* or *fmil-1* function. This finding suggests that additional PVD-expressed transcription factors and targets in our RNA-Seq data set could be required for the full complement of PVD→PVC synapses. Residual PVD→PVC synapses in *fmil-1* mutants could account for the robust initial response to optogenetic stimulation of PVD but are insufficient to sustain wild-type escape velocities during repetitive activation of the nociceptive circuit.

## Conclusions

Here we introduce a useful model in *C. elegans* for investigating the fundamental question of genetically encoded connectivity in the nervous system. The PVD nociceptive neuron synapses with target interneurons, PVC and AVA, to activate escape responses to noxious stimuli (*2*, *24*). In addition to its simplicity and well-defined behavioral output, the PVD circuit is generated in a discrete ventral cord region during larval development amenable to live cell imaging. We exploited these attributes to identify a conserved transcription factor, the LIM-homeodomain protein MEC-3, and its target, FMIL-1/Flamingo, a member of the CELSR subfamily of aGPCRs, as key determinants of PVD synapse formation with PVC and AVA. aGPCRs promote synaptogenesis across species and have been implicated in human neurological disorders, including bilateral frontoparietal polymicrogyria, spina bifida, and susceptibility to attention deficit hyperactivity disorder (*39*, *40*, *119–121*). Our work provides a genetically tractable experimental system for defining the molecular interactions and downstream signaling pathways whereby aGPCRs direct synapse formation between specific neurons, the fundamental developmental event that defines functional circuits in the brain.

Supplemental Table 1.

Strains used in this study.

Supplemental Table 2.

RNA-seq comparison between PVD and all cells.

Supplemental Table 3.

Sheet 1: PVD-enriched transcription factors.

Sheet 2: PVD-enriched transcription factors tested by RNAi for PVD→PVC synapse disruption.

Supplemental Table 4.

RNA-seq comparison between wild-type and *mec-3* PVD.

Supplemental Table 5.

Statistical comparisons for all experiments.

Supplemental Movie 1.

Appearance of PVD::myr::mCh in control animal. Playback rate is 7fps or 21min/sec.

Supplemental Movie 2.

Appearance of PVD::GFP::RAB-3 puncta in control animal. Playback rate is 7fps or 21min/sec.

Supplemental Movie 3.

Appearance of PVD::myr::mCh in *fmil-1* animal. Playback rate is 7fps or 21min/sec.

Supplemental Movie 4.

Appearance of PVD::GFP::RAB-3 puncta in *fmil-1* animal. Playback rate is 7fps or 21min/sec.

## Acknowledgements

We thank A. Brown, A. Gottschalk and S. Taylor for advice, O. Hobert and M. Van Hoven for reagents, E. Ozkan for unpublished results, and W. Thomson for DNA sequencing. Funded by NIH (1R01NS113559) to DMM, (F32NS117787) to TK, and HHMI funding to AD. Some strains were provided by NBRP, which is funded by the Japanese government. Some strains were provided by the CGC, which is funded by NIH Office of Research Infrastructure Programs (P40 OD010440). We also thank the Flow Cytometry Shared Resource for FACS support, supported by Ingram Cancer Center [P30 CA68485], DDRC [DK058404] and VANTAGE for sequencing support, supported by CTSA [5UL1 RR024975-03], Ingram Cancer Center [P30 CA68485], Vision Center [P30 EY08126], and NIH/NCRR [G20 RR030956].

---

**Supplemental Figure 1.**
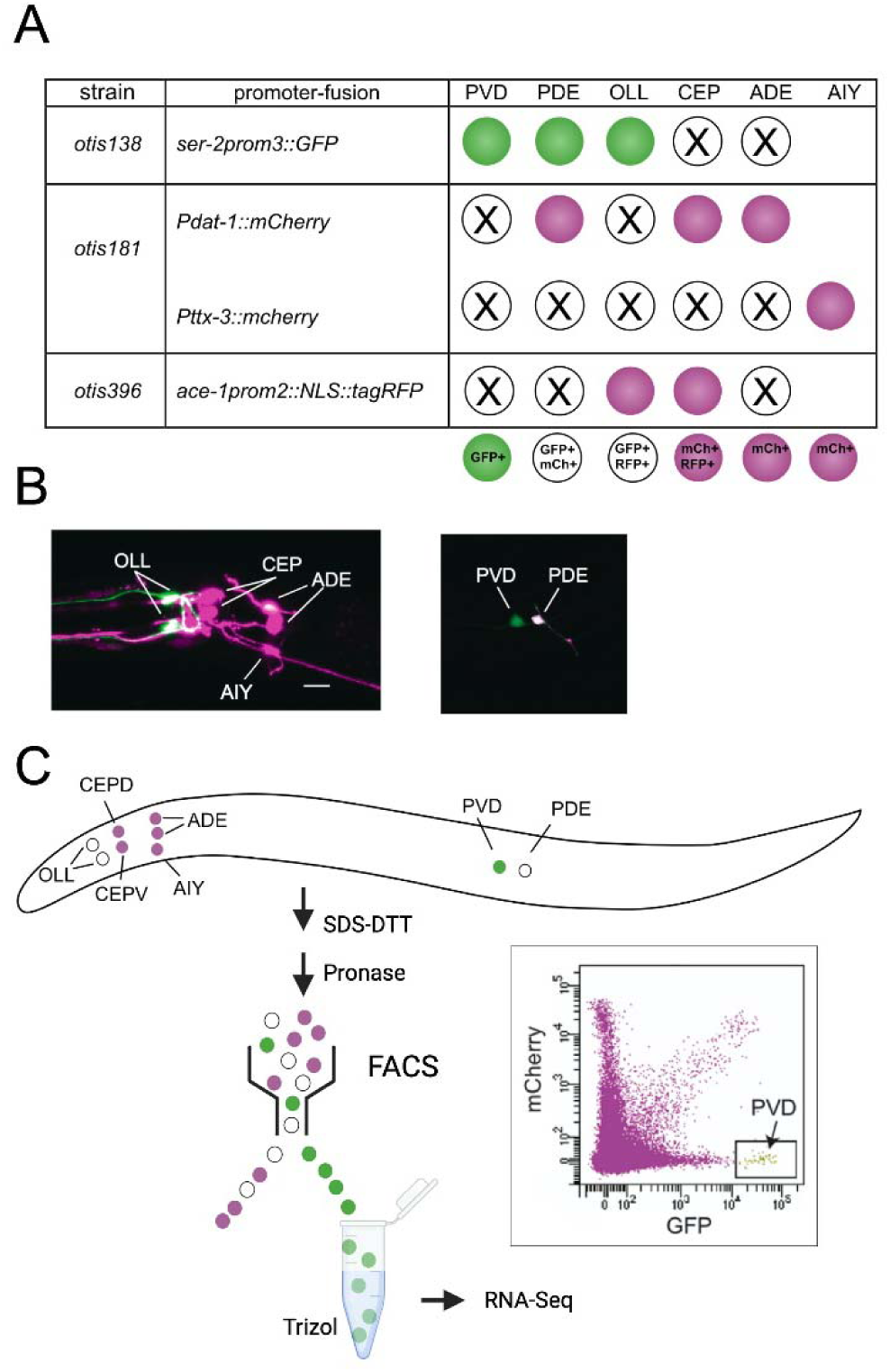
Transcriptional profiling of PVD nociceptive neurons to detect MEC-3-regulated genes. A. With intersectional labeling by three reporter strains (*ser2prom3::GFP, dat-1::RFP, ace-1prom2::NLS::tagRFP*), PVD is uniquely marked with GFP. B. Confocal images of head region (left) and posterior midbody (right) showing labeled neurons. Scale bar 10 μm. C. Cell dissociation for FACS isolation of PVD. Larvae were dissociated by successive treatments with SDS-DTT and pronase. FACS-isolated GFP+ neurons were collected in Trizol for RNA-isolation and sequencing.

**Supplemental Figure 2.**
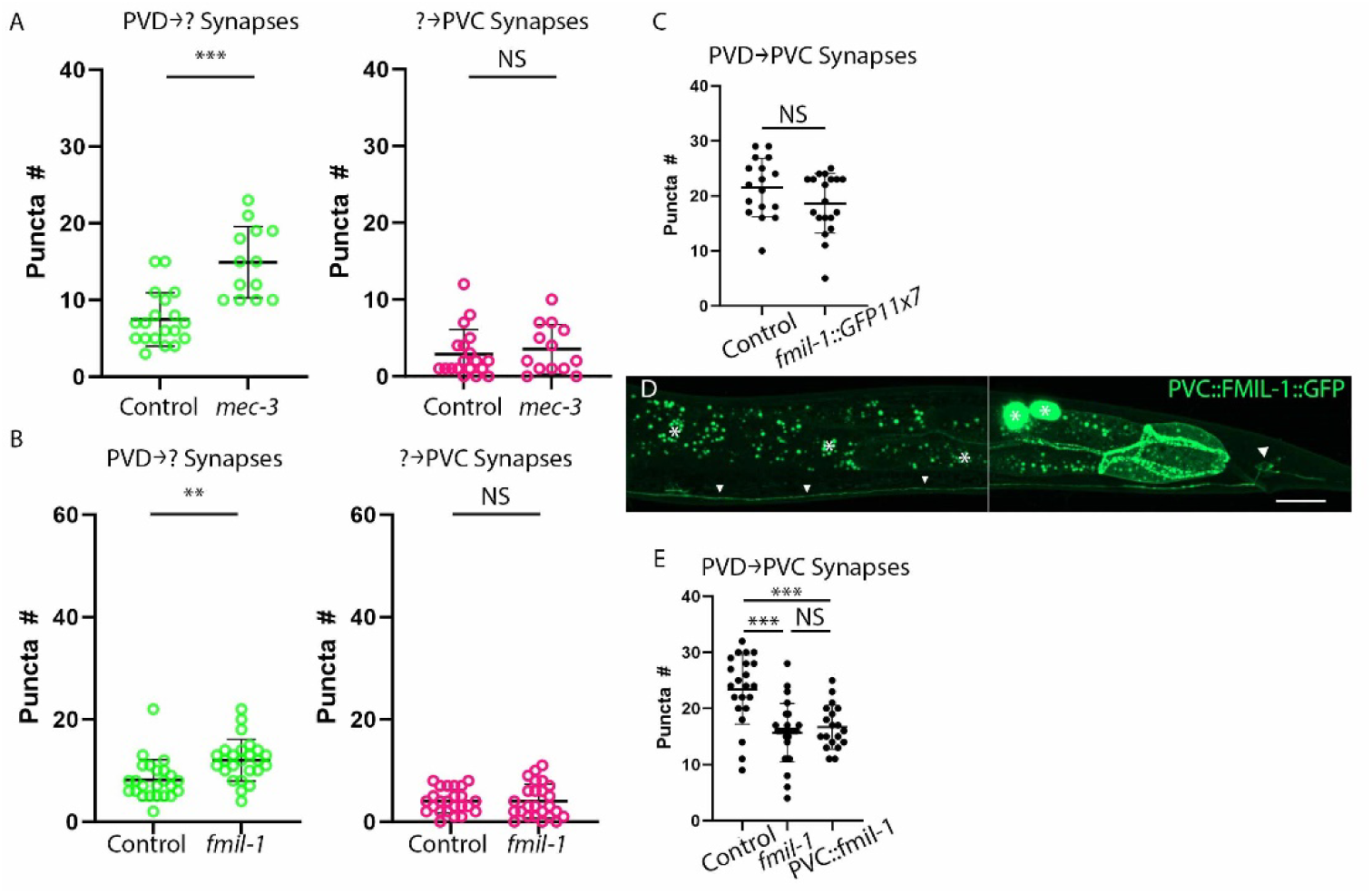
FMIL-1 acts pre-synaptically in PVD. A. Quantification of “unpaired” PVD pre-synapses and “unpaired” PVC post-synapses between control and *mec-3(gk1126)* nulls. Error bars are SD. ***p = 1.4x10^-5^, NS = Not Significant, Student’s t-test. B. Quantification of “unpaired” PVD pre-synapses and “unpaired” PVC post-synapses between control and *fmil-1(ok2115)* nulls. Error bars are SD. **p = 0.0015, NS = Not Significant, Student’s t-test. C. Endogenously tagged FMIL-1 does not disrupt PVD→PVC synapse formation. NS = Not Significant, Student’s t-test. D. PVC::FMIL-1::GFP expression. (Left) Midbody region with FMIL-1::GFP labeled PVC axon (arrowheads) in the ventral nerve cord. Asterisks denote gut autofluorescence. (Right) Posterior region showing PVC cell body (arrowhead). Asterisks denote the *unc-122::GFP* co-injection marker in coelomocytes. Scale bar 20µm. E. Quantification of PVD→PVC synapses shows PVC-specific expression of FMIL-1 (PVD::fmil-1) in *fmil-1(ok2115)* is not sufficient to restore PVD→PVC synapses to wild-type (Control) levels. ***p < 2.2x10^-4^, NS = Not Significant, one-way ANOVA with a Tukey’s post-hoc multiple comparison test.

**Supplemental Figure 3.**
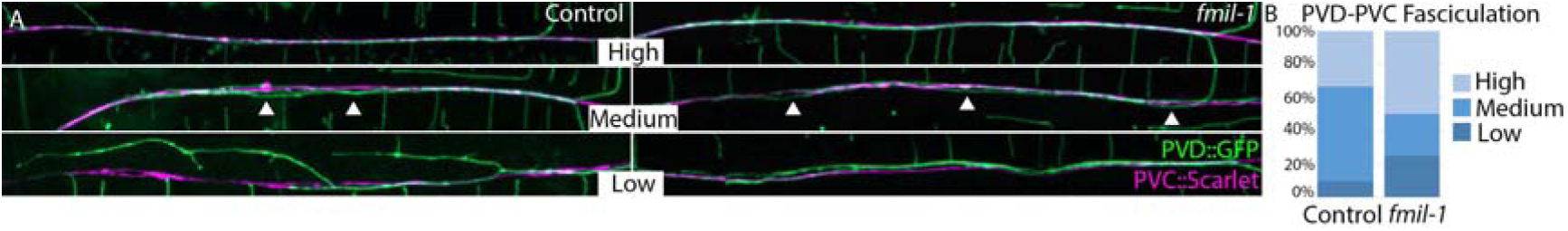
FMIL-1 is not required for PVD fasciculation with PVC in the ventral nerve cord. A. Representative images of high (top), medium (middle) and low (bottom) fasciculation between PVD (green) and PVC (magenta) in control (left) and *fmil-1* (right) animals. Arrowheads denote gaps between PVD and PVC. B. Quantification of high, medium and low fasciculation frequency in control and *fmil-1(ok2115)* animals.

**Supplemental Figure 4.**
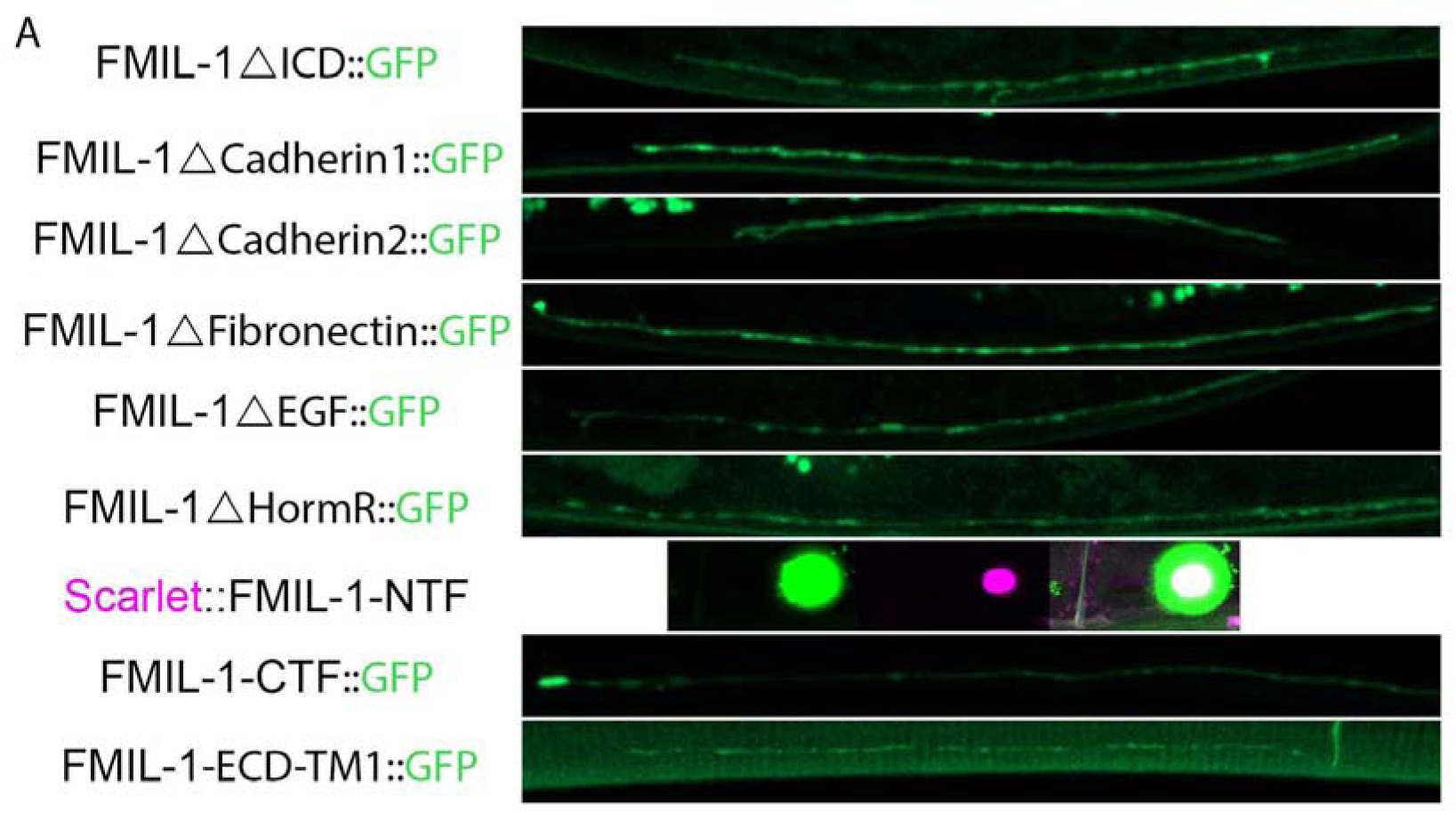
Alterations to FMIL-1 domains do not impact trafficking. A. Expression of FMIL-1::GFP ECD variant rescue constructs in the PVD axon. For Scarlet::FMIL-1-NTF, scarlet expression is shown in the center of a coelomocyte labeled with GFP, indicating that Scarlet::FMIL-1-NTF protein was secreted from PVD (*122*).

**Supplemental Figure 5.**
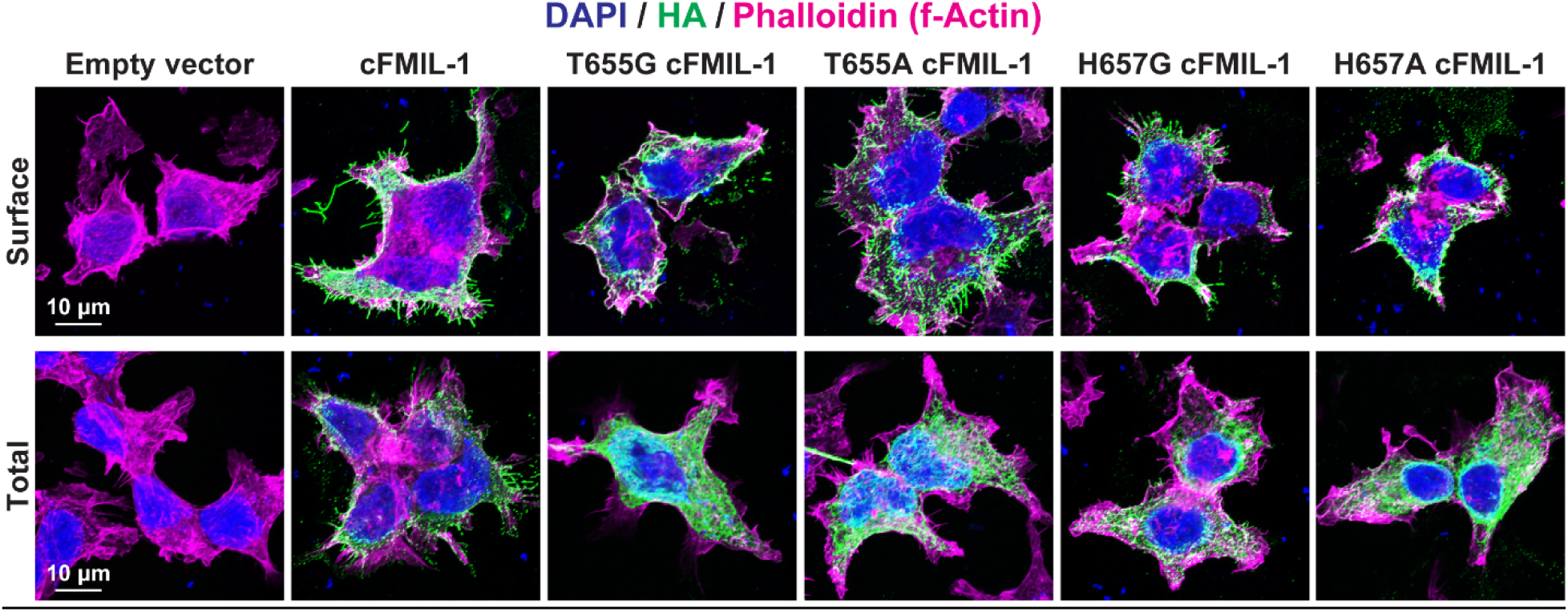
FMIL-1 Traffics to Surface of HEK293T cells. A. Representative images of surface expression levels of full-length cFMIL-1 relative to cFMIL-1 point mutants (T665G, T665A, H657G and H657A) in HEK293T cells. (Top) Cells were immunolabeled for surface HA in unpermeabilized conditions, followed by phalloidin (F-actin) and DAPI (nuclei). (Bottom) Cells were permeabilized and labeled for total HA together with phalloidin and DAPI.

**Supplemental Figure 6.**
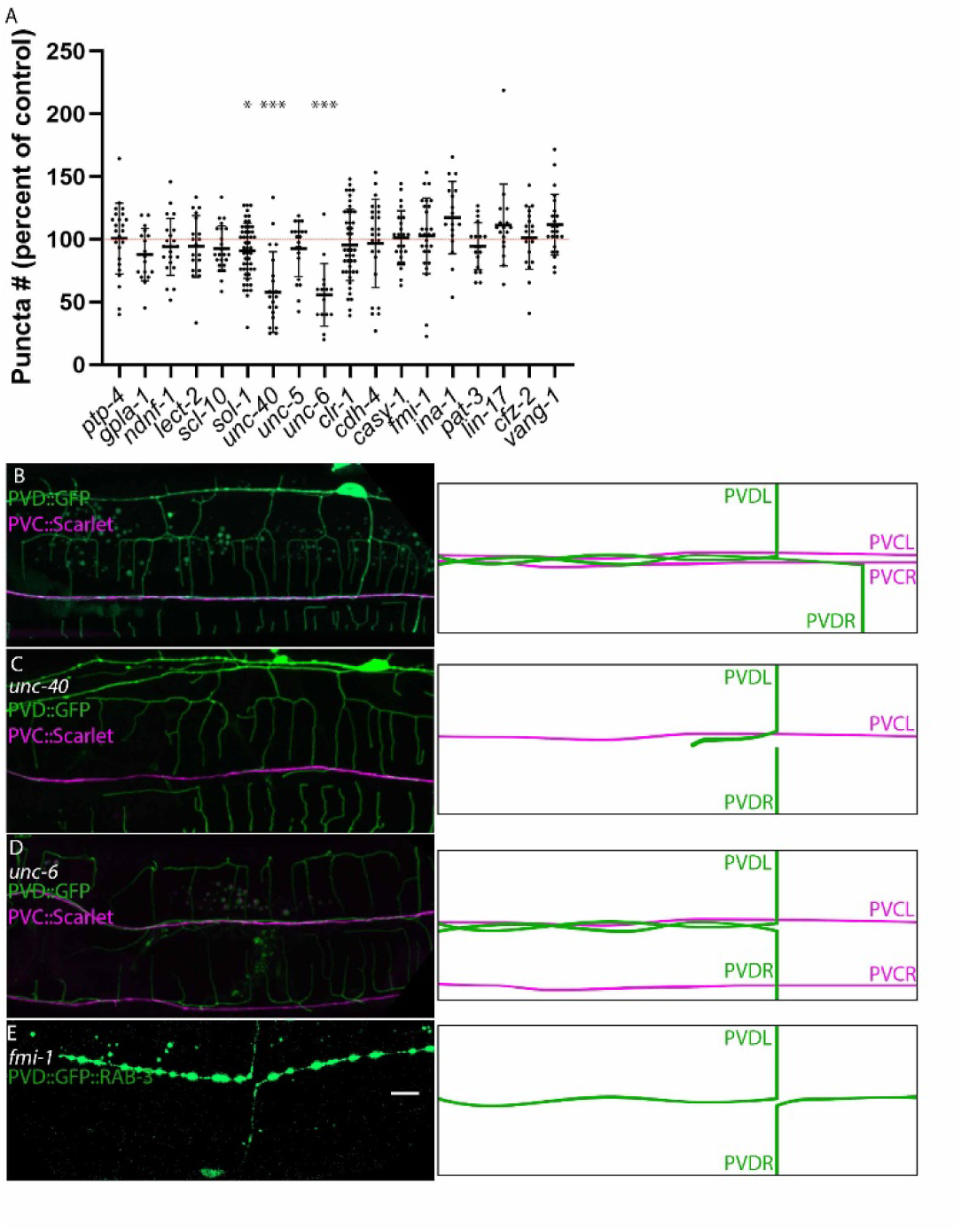
Potential FMIL-1 binding partners are not required for PVD→PVC synapses. A. Quantification of PVD→PVC synaptic puncta as a percent of paired wild-type control for candidate FMIL-1 interacting proteins. *p = 0.04, ***p < 2.1x10^-6^, Student’s t-test. B. (Left) PVD-PVC fasciculation in wild-type control. (Right) Schematic of PVD-PVC fasciculation. PVD dendrites not shown. C. (Left) Representative examples of defasciculated PVD vs PVC axons in *unc-40(e271)*; one PVD axon is missing. (Right) Schematic of PVD-PVC defasciculation defect in *unc-40*. PVD dendrites not shown. D. (Left) Representative examples of defasciculated PVD vs PVC axons in *unc-6(ev400)* mutants. In this example, the VNC (top of image) contains the PVD axons and one PVC process. The other PVC process is at the bottom of the image, likely in the lateral cord (arrowhead). (Right) Schematic of PVD-PVC defasciculation defect in *unc-6*. PVD dendrites not shown. E. (Left) In *fmi-1* mutants, the PVD axons occasionally turn posteriorly rather than anteriorly. Ventral view of PVD::GFP::RAB-3 highlights PVDL turning anteriorly and PVDR turning posteriorly. (Right) Schematic of PVD axonal defect. PVD dendrites not shown. Scale bar 10µm (applies to B-E).

