## Supplementary figures and images for "The Adhesion GPCR Flamingo-Like 1 (FMIL-1) Directs Synapse Formation in a Nociceptive Circuit"

### Supplemental figure 1

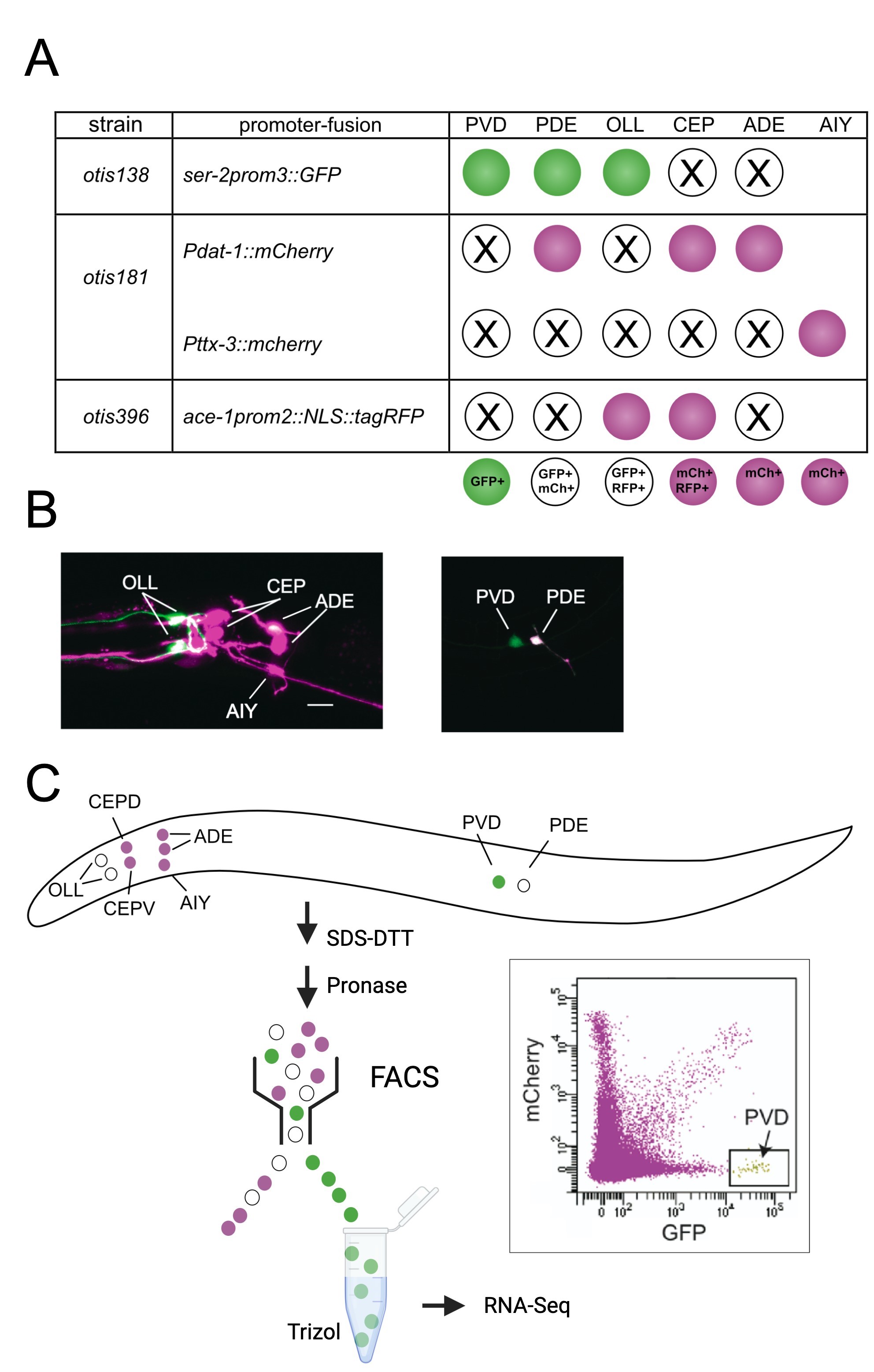

### Supplemental figure 2

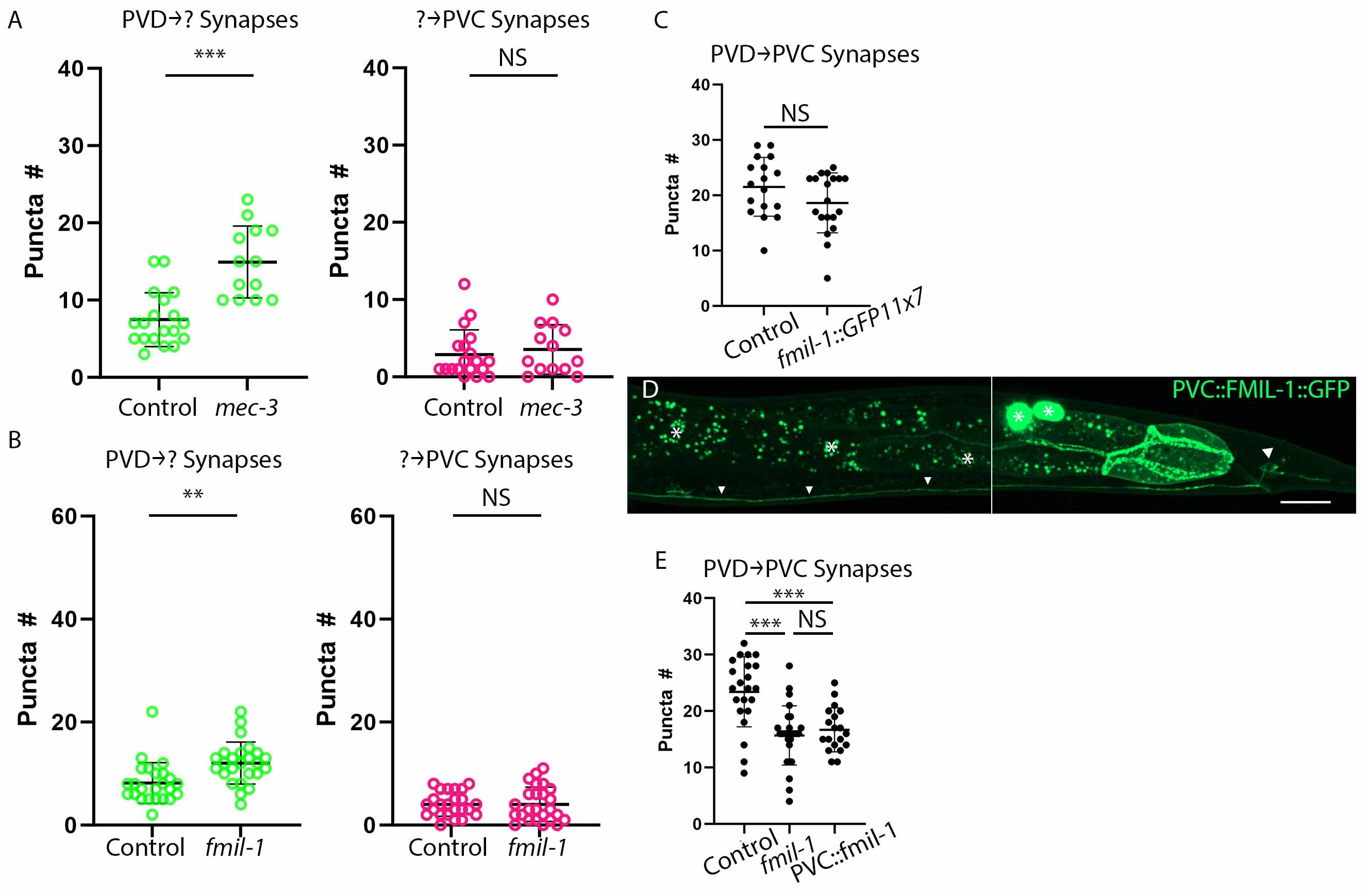

### Supplemental figure 3

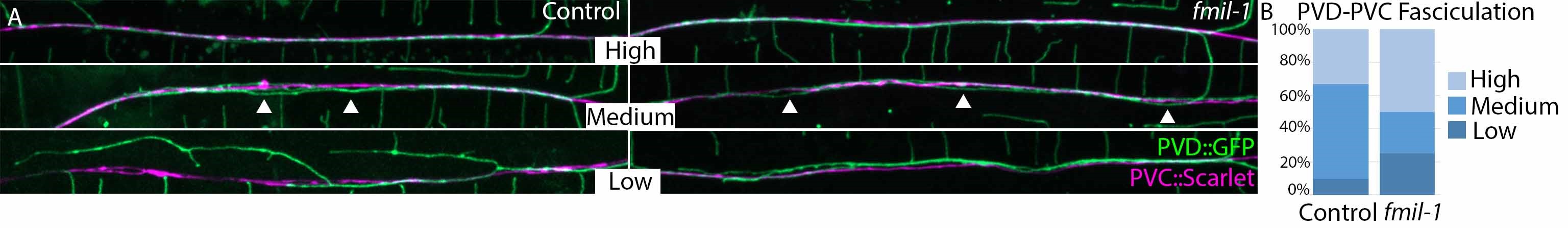

### Supplemental figure 4

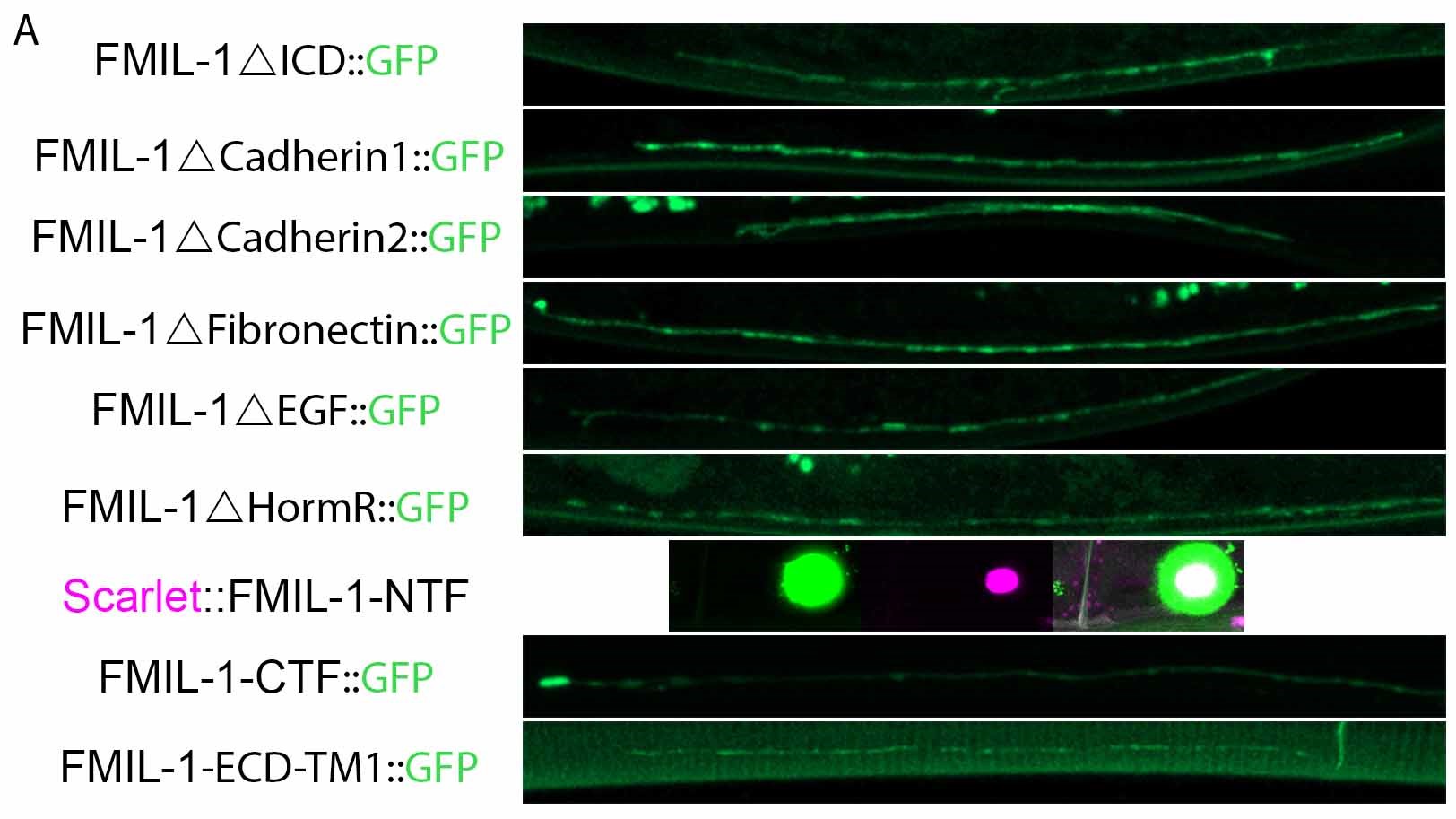

### Supplemental figure 5

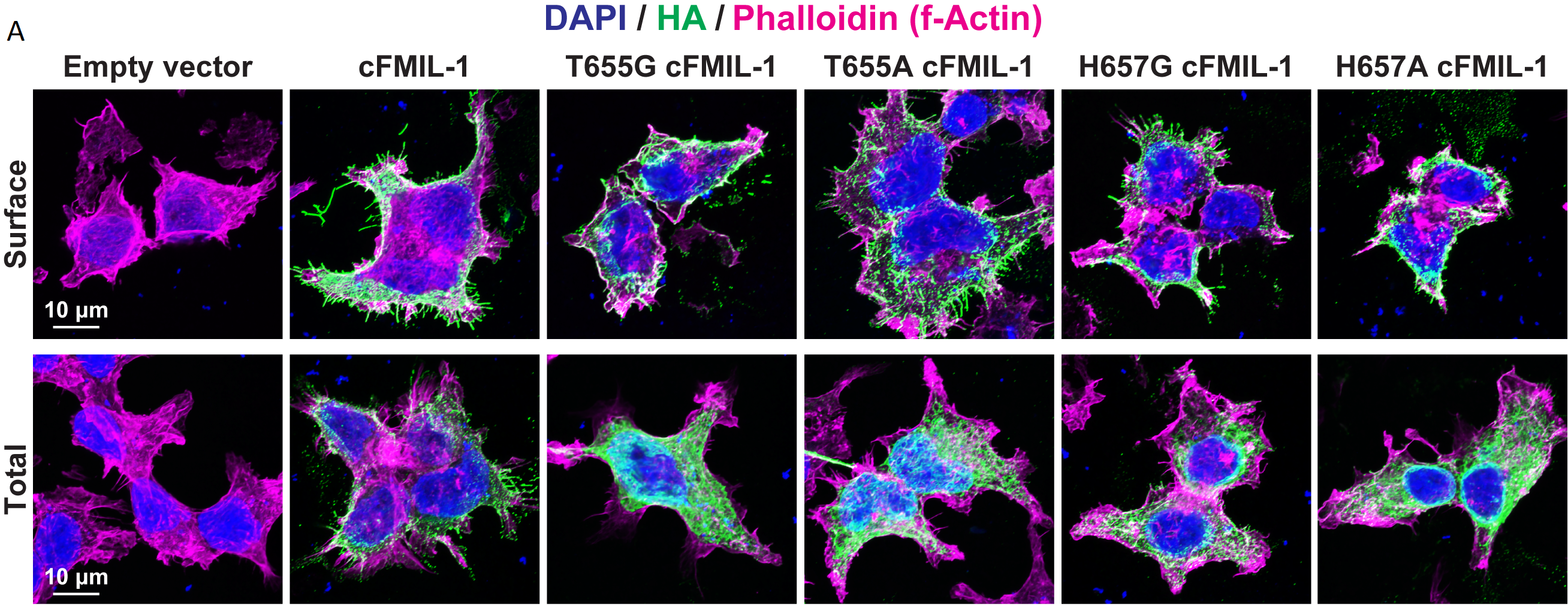

### Supplemental figure 6

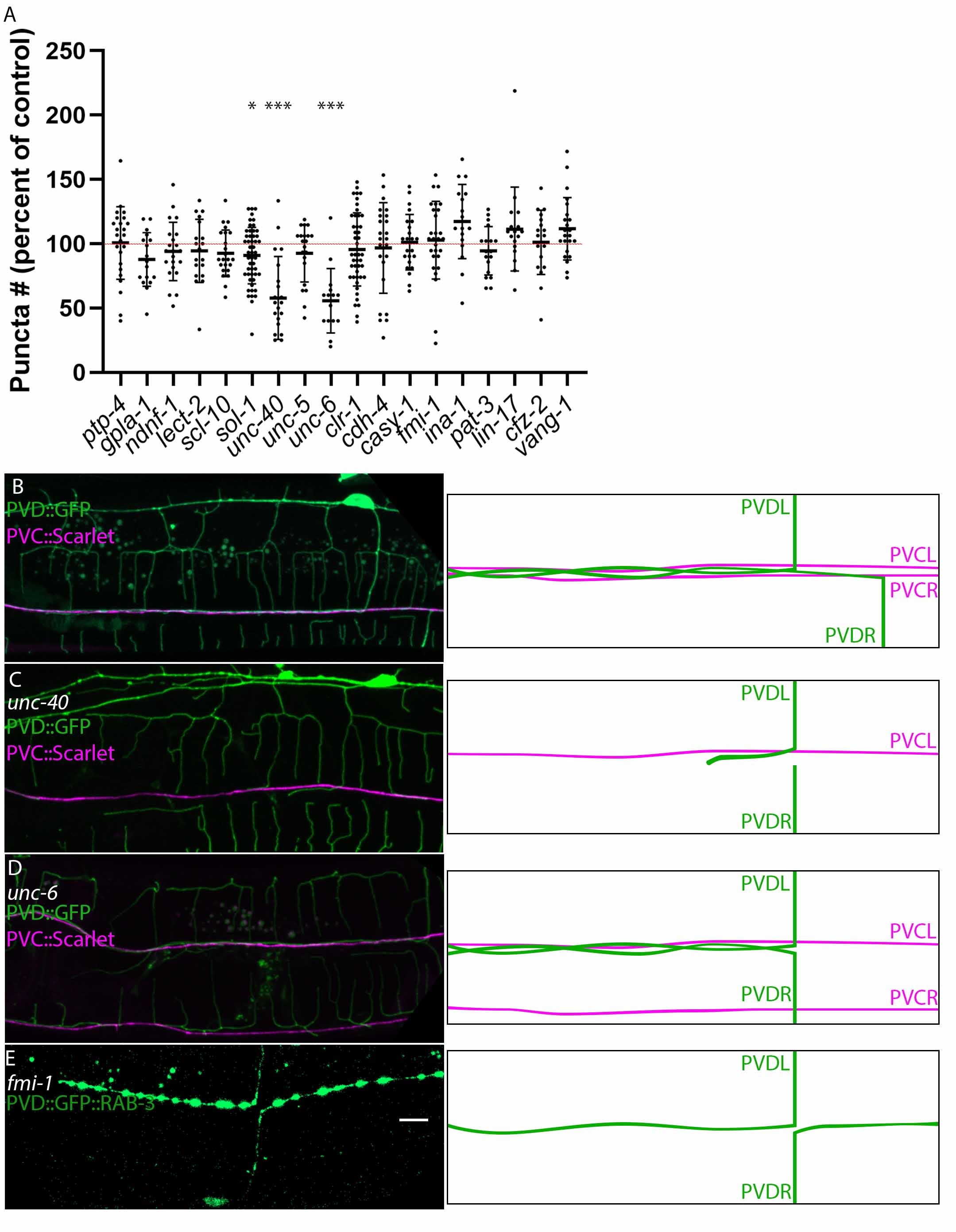
